# Structural snapshots of nitrosoglutathione binding and reactivity underlying S-nitrosylation of photosynthetic GAPDH

**DOI:** 10.1101/2022.05.03.490436

**Authors:** Edoardo Jun Mattioli, Jacopo Rossi, Maria Meloni, Marcello De Mia, Christophe H. Marchand, Andrea Tagliani, Silvia Fanti, Giuseppe Falini, Paolo Trost, Stéphane D. Lemaire, Simona Fermani, Matteo Calvaresi, Mirko Zaffagnini

## Abstract

S-nitrosylation is a redox post-translational modification widely recognized to play an important role in cellular signaling as it can modulate protein function and conformation. At the physiological level, nitrosoglutathione (GSNO) is considered the major physiological NO-releasing compound due to its ability to transfer the NO moiety to protein thiols. GSNO can also induce protein S-glutathionylation but the structural determinants regulating its redox specificity are not fully elucidated. In this study, we employed photosynthetic glyceraldehyde-3-phosphate dehydrogenase from *Chlamydomonas reinhardtii* (CrGAPA) to investigate the molecular mechanisms underlying GSNO-dependent thiol oxidation. We first observed that GSNO causes enzyme inhibition by specifically inducing S-nitrosylation. Treatment with reducing agents restores CrGAPA activity completely. While the cofactor NADP^+^ only partially protects from GSNO-mediated S-nitrosylation, the resultant inactivation is completely blocked by the presence of the substrate 1,3-bisphosphoglycerate, indicating that the S-nitrosylation of the catalytic Cys149 is responsible of CrGAPA inactivation. The crystal structures of CrGAPA in complex with NADP^+^ and NAD^+^ reveal a general structural similarity with other photosynthetic GAPDH. Starting from the 3D structure, we carried out molecular dynamics simulations to identify the protein residues involved in GSNO binding. Quantum mechanical/molecular mechanical calculations were performed to investigate the reaction mechanism of GSNO with CrGAPA Cys149 and to disclose the relative contribution of protein residues in modulating the activation barrier of the trans-nitrosylation reaction. Based on our findings, we provide functional and structural insights into the response of CrGAPA to GSNO-dependent regulation, possibly expanding the mechanistic features to other protein cysteines susceptible to be oxidatively modified by GSNO.

## Introduction

Protein redox post-translational modifications (PTMs) play an essential role in signaling pathways in most prokaryotes and eukaryotes, including photosynthetic organisms. Redox modifications mainly occur on the two sulfur-containing amino acids (*i.e.,* methionine and cysteine) due to their propensity to be oxidatively modified by reactive oxygen and nitrogen species (ROS and RNS, respectively) (1–4). While methionine-dependent regulation of plant processes is still an emerging field, protein cysteines are widely recognized to play a fundamental role in cell signaling, acting as regulatory molecular switches (5). To note, only cysteine residues that are found in the deprotonated state (*i.e.,* cysteine thiolates, -S^-^) are susceptible to oxidative modifications and various structural determinants contribute to the relative reactivity of the thiol group (5,6).

Nitric oxide (•NO, hereafter referred to as NO) is a relatively stable free radical recognized to act as a signaling molecule in controlling multiple physiological processes in both animal and plant systems (7,8). The biological effects of NO are thought to be primarily linked to a redox PTM named S-nitrosylation (also referred to as protein S-nitrosation) (5,9). This oxidative modification consists in the formation of a nitrosothiol (-SNO) between NO and a redox-reactive protein cysteine and results in the alteration of enzyme activities, protein conformation and stability, as well as interactions with other macromolecules including proteins and nucleic acids (4). The formation of nitrosothiols can occur through the direct reaction of NO with thiyl radical (-S•), or they can derive from the addition of a nitrosonium group (NO^+^) to a cysteine thiolate (4). At the physiological level, dinitrogen trioxide (N_2_O_3_) and nitrosoglutathione (GSNO) are considered the prominent nitrosylating agents due to their ability to donate their NO^+^ moiety to a target cysteine (10). While the reaction of N_2_O_3_ with cysteine residues does not seem to require specific structural constraints, the interaction of GSNO with target cysteines is supposed to be assisted by the presence of acidic and basic residues flanking or surrounding the protein thiol in the primary or tertiary sequence, respectively (11,12). This structural feature has been named GSNO binding motif and it functions to both enhance proton release from the cysteine thiol and ensure a proper binding of the nitrosylating agent (13). The identification of a -SNO consensus motif in target proteins has been sought, but a universal pattern has not been established yet. Besides acting as a trans-nitrosylating agent, GSNO can also induce S-glutathionylation, a redox modification consisting in the formation of a mixed disulfide between a molecule of glutathione and a protein cysteine (14–16). S-glutathionylation shares with S-nitrosylation the ability to regulate protein function and conformation, and it can be induced by alternative mechanisms involving a thiol/disulfide exchange mediated by oxidized glutathione (GSSG) or H_2_O_2_-dependent primary oxidation of a cystine thiol to sulfenic acid followed by the spontaneous reaction with reduced glutathione (GSH) (6).

In the last decades, proteomic-based approaches identified hundreds of proteins undergoing S-nitrosylation in plants ((5) and references therein), highlighting the importance of this redox modification in the control of multiple cellular processes such as pathogen resistance, immune response, and carbon-related metabolic pathways (4, 5, 17–21). Notwithstanding the numerous putative targets, glyceraldehyde-3-phosphate dehydrogenase (GAPDH) has been found as a prominent target and widely used to study the molecular mechanisms underlying NO-dependent thiol modifications (22–26). In plants, GAPDH comprises several isoforms participating in the glycolytic pathway in the cytoplasm and the stroma (NAD(H)-dependent GAPC and GAPCp isoforms, respectively) and in the Calvin-Benson-Bassham (CBB) cycle (*i.e.,* the reductive pentose phosphate cycle) in the stroma (NADP(H)-dependent GAPA and GAPA/B isozymes). Regardless of their metabolic function, the catalytic mechanism of GAPDH enzymes strictly depends on a reactive cysteine located in the active site (22). The reactivity of this residue (hereafter numbered as Cys149, (27)) is crucial for the nucleophilic attack on the substrate and it is fostered by an interaction with the proximal His176 that attracts the proton from the sulfur atom stabilizing the thiolate state (-S^−^) (22). Besides being required for the catalysis, the deprotonation of Cys149 makes it sensitive to several types of redox modifications (*e.g*., S-nitrosylation, S-glutathionylation, persulfidation, and sulfenic acid formation), which can unavoidably alter its functionality (23,24,27–31). In previous studies, glycolytic GAPCs from *Arabidopsis thaliana* (AtGAPCs) were employed to investigate the molecular mechanisms controlling S-nitrosylation/denitrosylation reactions (23,24). Notably, AtGAPC activity is modulated by reversible S-nitrosylation of its catalytic cysteine and S-nitrosothiol formation is induced by GSNO-dependent transnitrosylation and controlled by GSH/GSNO ratios. Besides forming a nitrosothiol, GSNO also induced S-glutathionylation of AtGAPC even though to a minor extent (23). Activity modulation by GSNO was also observed for GAPDH-related activities when assayed in Arabidopsis protein extracts (25), and for the recombinant form of photosynthetic GAPDH from *Chlamydomonas reinhardtii* (26).

Considering the prominent role of GSNO as a mediator of NO-dependent biological activities and the thiol-dependent regulatory switch of GAPDH activity, we sought to elucidate the structural determinants that control GSNO binding and reactivity as well as the molecular mechanisms underlying the GSNO-dependent oxidation of plant GAPDH. To this aim, we employed a combination of biochemical, structural, and computational approaches to investigate the regulatory role of GSNO on GAPA from *Chlamydomonas reinhardtii* (CrGAPA). GAPA is, the unique photosynthetic GAPDH isoform present in green algae. Exposure to GSNO led to CrGAPA inactivation via specific S-nitrosylation of its catalytic cysteine. The binding of CrGAPA substrate (*i.e.*, 1-3-bisphosphoglycerate, BPGA) fully protects the enzyme, while the cofactor NADP^+^ causes a partial protection delaying the inactivation kinetics. Determination of the crystal structures of CrGAPA bound to both NAD^+^ and NADP^+^ allowed the comparison with other structurally known plant GAPDH, and it was instrumental to establish the protein residues involved in the GSNO binding using molecular dynamics (MD). The reaction between GSNO and the catalytic cysteine was investigated using a quantum mechanical/molecular mechanical (QM/MM) approach revealing mechanistic features of GSNO reactivity with CrGAPA catalytic cysteine. Based on our findings, we provide mechanistic insights into the response of a photosynthetic GAPDH to GSNO-dependent regulation, possibly extending this analysis to cysteine microenvironments from other proteins that are susceptible to be oxidatively modified by GSNO.

## Material and Methods

### Chemicals and enzymes

N-[6-(Biotinamido)hexyl]-3’-(2’-pyridyldithio)proprionamide (HPDP-biotin) was purchased from Pierce Biotechnology. All chemicals and enzymes were obtained from Merck Life Science unless otherwise specified. Recombinant CrGAPA was expressed and purified according to (26). The concentration of purified CrGAPA was determined spectrophotometrically using a molar extinction coefficient at 280 nm (*ε*_280_) of 36,565 M^-1^ cm^-1^.

### Crystallization and data collection

Purified CrGapA was concentrated to 10 mg/ml in 30 mM Tris-HCl, 1 mM EDTA (pH 7.9), and 1 mM NAD^+^ or NADP^+^ and crystallized by the hanging drop vapor-diffusion method at 293 K. Protein solution aliquots of 2 μl were mixed to an equal volume of reservoir and the final drop was equilibrated against 750 μl reservoir solution. Aggregate crystals appeared in 10-15 days using 1.8-2.0 M (NH_4_)_2_SO_4_ as precipitant and 0.1 M Tris-HCl pH 7.5-8.5 or Hepes-NaOH pH 7.5, thus the conditions were optimized decreasing the precipitant concentration or protein concentration or both. Best crystals used for further diffraction experiments, grew with a protein concentration ranging from 5 to 10 mg/ml, 1.2-1.6 M (NH_4_)_2_SO_4_, 0.1 M Tris-HCl pH 7.5-8.5 or 0.1 M Bicine pH 9.5 (only for oxidized NADP^+^-CrGapA). Crystals were harvest by a cryo-loop, briefly soaked in the cryo-protectant solution (1.6 M (NH_4_)_2_SO_4_, 20% glycerol, and 2 mM NAD^+^ or NADP^+^) and finally frozen in liquid nitrogen.

Diffraction data were collected at the Elettra synchrotron radiation source (Trieste, beam line XRD1) at 100 K using a wavelength of 1.0 Å, an oscillation angle (ΔΦ) of 0.5° for NAD^+^- and oxidized NADP^+^-CrGapA and 0.3° for NADP^+^-CrGapA, and a sample-to-detector (Pilatus 2M) distance (d) of 160, 190 and 200 mm for NADP^+^-, oxidized NADP^+^- and NAD^+^-CrGapA, respectively. Data were processed using XDS (32) and scaled with AIMLESS (33). The correct space group was determined with POINTLESS (34) and confirmed in the structure solution stage. Unit cell parameters and statistics are reported in Supplemental Table 1.

### Structure solution and refinement

CrGapA structures were solved by molecular replacement using the program MOLREP (35) from CCP4 package (36), using the structure of SoGAPA (PDB ID code: 1JN0; (37)) excluding non-proteins atoms and water molecules, as a search model. The refinement was performed with REFMAC5 7.1.004 (38) from CCP4 package (36), selecting 5% of reflection for R_free_ calculation. The molecular graphic software COOT (39) was used for manual rebuilding and modelling of the missing atoms in the electron density map and to add solvent molecules. Water molecules were automatically added and, after a visual inspection, confirmed in the model if the relative electron density value in the (2F_O_ – F_C_) maps exceeded 0.19 e-Å-3 (1.0 σ) and if they fell into an appropriate hydrogen bonding environment. Inspection of the Fourier difference maps of CrGapA crystals clearly showed additional electron densities attributed to the cofactors (NAD^+^ or NADP^+^) and to an oxidized thiol group (-SO3) of the catalytic Cys149. For NADP^+^-CrGapA structures the last refinement cycle was performed with PHENIX (40). Final refinement statistics are reported in Supplemental Table 1.

The superpositions among structures have been performed by LSQKAB (41) from CCP4 package (36). The structures have been validated using MolProbity (42). Figures were generated using Pymol (The PyMOL Molecular Graphics System, Schrödinger, LLC).

### Activity assay

CrGAPA activity was monitored as described previously (29,43). Briefly, the reaction was measured spectrophotometrically at 340 nm and 25 °C in an assay mixture containing 50 mM Tris-HCl (pH 7.5), 1 mM EDTA, 5 mM MgCl2, 3 mM 3-phosphoglycerate, 5 units/ml of yeast 3-phosphoglycerate kinase, 2 mM ATP, and 0.2 mM NADPH.

### Treatment of CrGAPA with GSNO

CrGAPA (2 μM) was incubated in 50 mM Tris-HCl buffer (pH 7.9) in the presence of different concentrations of GSNO. After 30 min incubation, an aliquot of the sample (5 μl) was withdrawn for the assay of enzyme activity. Substrate protection was performed by pre-incubating (5 min) the protein in the presence of a 1,3-bisphosphoglycerate-generating system (3 mM 3-phosphoglycerate, 5 units/ml of 3-phosphoglycerate kinase, and 2 mM ATP) or in the presence of 0.2 mM NADP^+^. The reversibility of GSNO-mediated CrGAPA inactivation was assessed by measuring protein activity after incubation for 10 min in the presence of 20 mM dithiothreitol (DTT). The S-nitrosylation signal of GSNO-treated CrGAPA was assessed using the biotin switch technique as described in (23).

### MALDI–TOF (matrix-assisted laser-desorption ionization–time-of-flight) mass spectrometry

CrGAPA was treated for 30 min with 1 mM GSNO and MALDI-TOF mass spectrometry analysis was performed before and after incubation with 20 mM DTT for 30 min. The samples were analyzed as described in (31,44).

### Molecular dynamics (MD) simulations

#### Setting the MD simulation

MD simulations were performed using the AMBER 16 package (45). The FF14SB force field (46) was used to model CrGAPA, GSNO and GS^−^. For the nitrosylated cysteine an *ad hoc* force field developed by Han (47) was used. NADP^+^ was modelled with force field parameters calculated by (48). The charges of GSNO and GS^−^ were determined using the Merz-Singh-Kollman scheme (49). All simulations were performed with explicit solvent by using the TIP3P water model (50).

#### Minimization, equilibration and MD production

500 steps of steepest descent minimization, followed by additional 9500 steps of conjugate gradient minimization were performed with PMEMD (45). The minimized structure was used as starting point for the equilibration process. Particle Mesh Ewald summation was used throughout (with cut off radius of 10.0 Å), H-atoms were considered by the SHAKE algorithm and a time step of 2 fs was applied in all MD runs. 1ns of heating to 298 K within an NPT ensemble and temperature coupling according to Andersen was used to equilibrate the system. A MD trajectory of 100 ns is then produced. Snapshot structures were saved into individual trajectory files every 1000-time steps, *i.e.*, every 2 ps of molecular dynamics.

#### Post Processing of Trajectories

Trajectories obtained from MD simulations were post-processed using CPPTRAJ. (50,51) 1000 snapshots were extracted from the calculated trajectory (1 snapshot each 100 ps) to estimate the contributions to the binding free energy of the single amino acids of CrGAPA with GSNO and GS^-^, using the MM/GBSA approach (52).

### QM/MM calculations

#### Determination of the potential energy surface (PES) of the transnitrosylation reaction

QM/MM calculations were carried out according to ONIOM scheme (53) as implemented in Gaussian16 (54). The inner QM layer consists in the reacting part of the system, *i.e.*, H_3_CS^-^ + CH_3_SNO and was described at the DFT level using the M06-2X functional (55) and the 6-311 + +G** basis set (54). The outer layer was described at the molecular mechanics (MM) level employing the parameters used in the MD simulations. The structure of the various critical points (minima and saddle points) was fully optimized. Frequency calculations were carried out at the same level of theory to check the nature of critical points and, to calculate Gibbs free energies. Water solvation was modelled using PCM model as implemented in Gaussian (55).

#### Fingerprint analysis

To quantify the catalytic effect of the residues surrounding the cysteines (within 5 Å) involved in the transnitrosylation process we recomputed the activation energy of the transnitrosylation reaction, calculating the electrostatic (Coulomb) effect of the *i*th residue on the QM region in the reactant and in the transition state (fingerprint analysis) (56–58). The analyses demonstrate the stabilizing/destabilizing effects exerted by the various residues.

### Accession Numbers

The atomic coordinates and structure factors of CrGapA structures have been deposited in the Protein Data Bank with the accession codes: 7ZQ3, 7ZQK, and 7ZQ4 for NADP^+^-, NAD^+^- and oxidized NADP^+^-CrGapA, respectively.

## Results

### Comparing CrGAPA sequence with plastidial GAPDHs from photosynthetic organisms

Multiple sequence alignments reveal that CrGAPA shows a relatively high similarity with photosynthetic GAPDH from land plants and microalgae (76-80% sequence identity; Supplemental Figures 1 and 2), while the sequence identity slightly decreases (64-68%) when we compared CrGAPA with homologs from cyanobacterial species (Supplemental Figures 3). Among photosynthetic GAPDH isoforms, the catalytic dyad Cys149/His176 and the majority of residues participating in the stabilization of the cofactors NADP(H) and NAD(H) and specificity towards NADP(H) are fully conserved (see below and Supplemental Figures 1-3). Multiple alignment of primary sequences was also instrumental to assess Cys conservation in photosynthetic GAPDH. CrGAPA shows in its primary structure four cysteines (Cys) at position 18, 149, 153, and 285 (Supplemental Figure 1). Cys18 and Cys149 are strictly conserved in photosynthetic GAPA isozymes, while Cys153 and Cys285 are absent in two microalgal species (*i.e., Ostreococcus tauri* and *Micromonas pusilla)* and cyanobacterial enzymes (*i.e., Synechococcus elongatus* PCC7942 and *Thermosynechococcus elongatus*), respectively, and are both replaced by glycine residues (Supplemental Figures 1-3).

### Three-dimensional structure of CrGAPA and structural comparison with photosynthetic GAPDH isoforms

In order to determine the structural features of CrGAPA, the enzyme was expressed in *E. coli* and purified to homogeneity by metal affinity chromatography. The recombinant protein contains 349 amino acids (mature protein plus the MHHHHHHM peptide) with a calculated molecular weight of 38103.8 Da consistent with SDS-PAGE analysis (Supplemental Figure 4). The crystal structures of CrGAPA complexed with both cofactors NADP^+^ and NAD^+^ (NADP- and NAD-CrGAPA) have been determined at a resolution of 1.5 and 2.2 Å, respectively (Supplemental Table 1). An additional structure of the enzyme complexed with NADP^+^ and showing the catalytic Cys149 oxidized to sulphinate/sulphonate (-SO_2_^−^/-SO_3_^−^) is also reported (Supplemental Table 1). All crystals are isomorphous and their asymmetric unit contains a dimer (chains named O and R) generating the whole tetramer by a crystallographic 2-fold axis corresponding to a molecular symmetry axis (Figure 1). The superimposition of the two independent chains determines a root mean square deviation (rmsd) of 0.21 Å (326 aligned C_α_ atoms) and 0.30 Å (335 aligned C_α_ atoms) in the case of NADP- and NAD-CrGapA, respectively. In reduced and oxidized NADP-CrGAPA structure, the well-defined electron density allowed the building of the whole non-cleavable His-tag (MHHHHHHM) at the N-terminal end of chain R. This portion disordered in chain O, breaks the 222 symmetry of the tetrameric structure which became a dimer of dimers (C2 symmetry) and is stabilized by interactions with symmetry related molecules (Supplemental Figure 5A, B).

**Figure 1.**
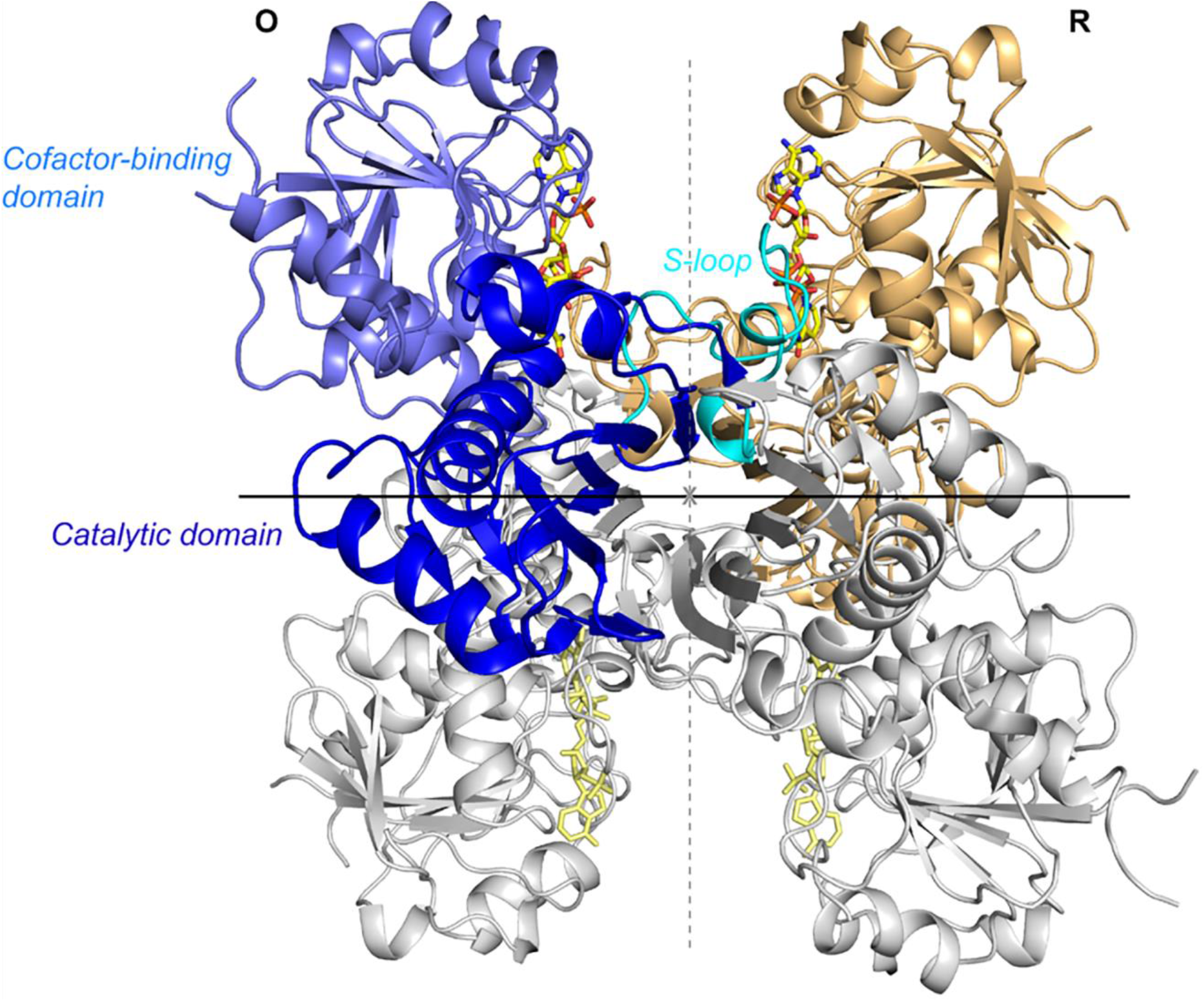
Crystal structure of CrGAPA tetramer. Ribbon representation of the CrGAPA tetramer. The dimer OR corresponds to the asymmetric unit, while the dimer generated by the 2-fold crystallographic axis, is reported in gray. The crystallographic symmetry axis is coincident with one of the symmetry molecular axis (in black), the other two symmetry molecular axes are represented with dashed lines (gray). In chain O, the different domains are highlighted: cofactor-binding domain in light blue, catalytic domain in blue and S-loop in cyan. The cofactor (NADP^+^) bound to each monomer, is represented in stick.

Besides CrGAPA, the crystal structures of photosynthetic GAPDH isoforms have been determined for spinach and Arabidopsis enzymes (SoGAPA and AtGAPA, respectively) (37,59–61) and for homologs from two cyanobacteria (*Synechococcus elongatus* PCC7942 and *Thermosynechococcus elongatus;* (62–65). As expected from the high sequence homology (64-80%), the 3D structure of photosynthetic GAPDH is highly conserved. The superimpositions of CrGAPA crystal structures with those of SoGAPA and AtGAPA give an average rmsd of 0.43-0.56 Å for monomers and 0.87 Å for tetramers. Structural conservation is also observed when we compared CrGAPA with cyanobacterial NADP(H)-dependent GAPDH, displaying rmsd ranging from 0.55 to 0.77 Å for monomer and from 0.84 to 1.18 Å for tetramer superimpositions.

### Domain organization, cofactor-binding, and catalytic sites of CrGAPA

Like in other GAPDH (37,61,66), each CrGAPA monomer consists of a cofactor-binding domain and a catalytic domain. The first one comprises residues 1-147 and 313-334 and shows the structurally conserved Rossmann fold motif typical of enzymes using nucleotide cofactors and an additional antiparallel β-sheet (Figure 1). The catalytic domain stretching from residues 148 to 312, is composed by a seven-stranded mixed β-sheets, three α-helices, and an ordered loop named S-loop (residues 177 to 203) which forms the interface with the adjacent subunit and contributes to the set-in place and binding of the cofactor (Figure 1).

Based on the electron density, we recognized that each CrGAPA monomer contains the coenzyme (NADP^+^ or NAD^+^) bound in an extended conformation through hydrogen bonds and electrostatic interactions with protein residues and water molecules (Figure 2A). The adenine and nicotinamide rings are roughly perpendicular to the average planes of the neighboring riboses. The first one is sandwiched between the methyl group of Thr96 and the guanidium group of Arg77 in the NADP-bound structure or the hydroxyl group of Ser33 in the NAD-bound structure (Figures 2A-C). The nicotinamide ring orientation is determined by an intramolecular hydrogen bond between the N7 and the O1 of the nicotinamide moiety (NO1-NN7 = 2.9 Å in both subunits of NADP-CrGAPA and 3.1 and 2.8 Å in O and R subunits of NAD-CrGAPA), and hydrophobic interactions with side chains of the strictly conserved Ile11 and Tyr317 (Figure 2A). The backbone nitrogen atoms of Gly9, Arg10, and Ile11 are involved in the stabilization of the central phosphate groups. The 2’-phosphate group in the adenine ribose of NADP^+^ forms a salt-bridge with the Arg77 and hydrogen bonds with Ser33 and Ser188 of the adjacent subunit, and various water molecules (Figure 2B). When NAD^+^ binds to the enzyme the hydroxyl groups of the adenine ribose form hydrogen bonds with water molecules and uniquely in chain R, with Ser188 of the adjacent subunit O (Figure 2C). Unlike other photosynthetic GAPDH (*i.e.*, GAPA from spinach, Arabidopsis, and the cyanobacterium *Thermosynechococcus elongatus*), the highly conserved Asp32 (Supplemental Figures 1-3), which is involved in the stabilization of NAD(H) (60,61,65), does not participate in the cofactor binding in CrGAPA lying at more than 4.5 Å from the hydroxyl groups of the adenine ribose (Figure 2C). The replacement of the catalytically preferred cofactor NADP^+^ with NAD^+^ does not significantly alter either the monomer or the tetramer folding. Indeed, the superimpositions of monomers and tetramers give a rmsd of 0.23-0.36 Å (333 aligned C_α_ atoms) and 0.32 Å (1326 aligned C_α_ atoms), respectively. Even the protein portion 31-36, which was observed to undergo conformational changes depending on the bound cofactor in photosynthetic GAPA from spinach (60), perfectly superimposes in NADP- and NAD-CrGAPA structures.

**Figure 2.**
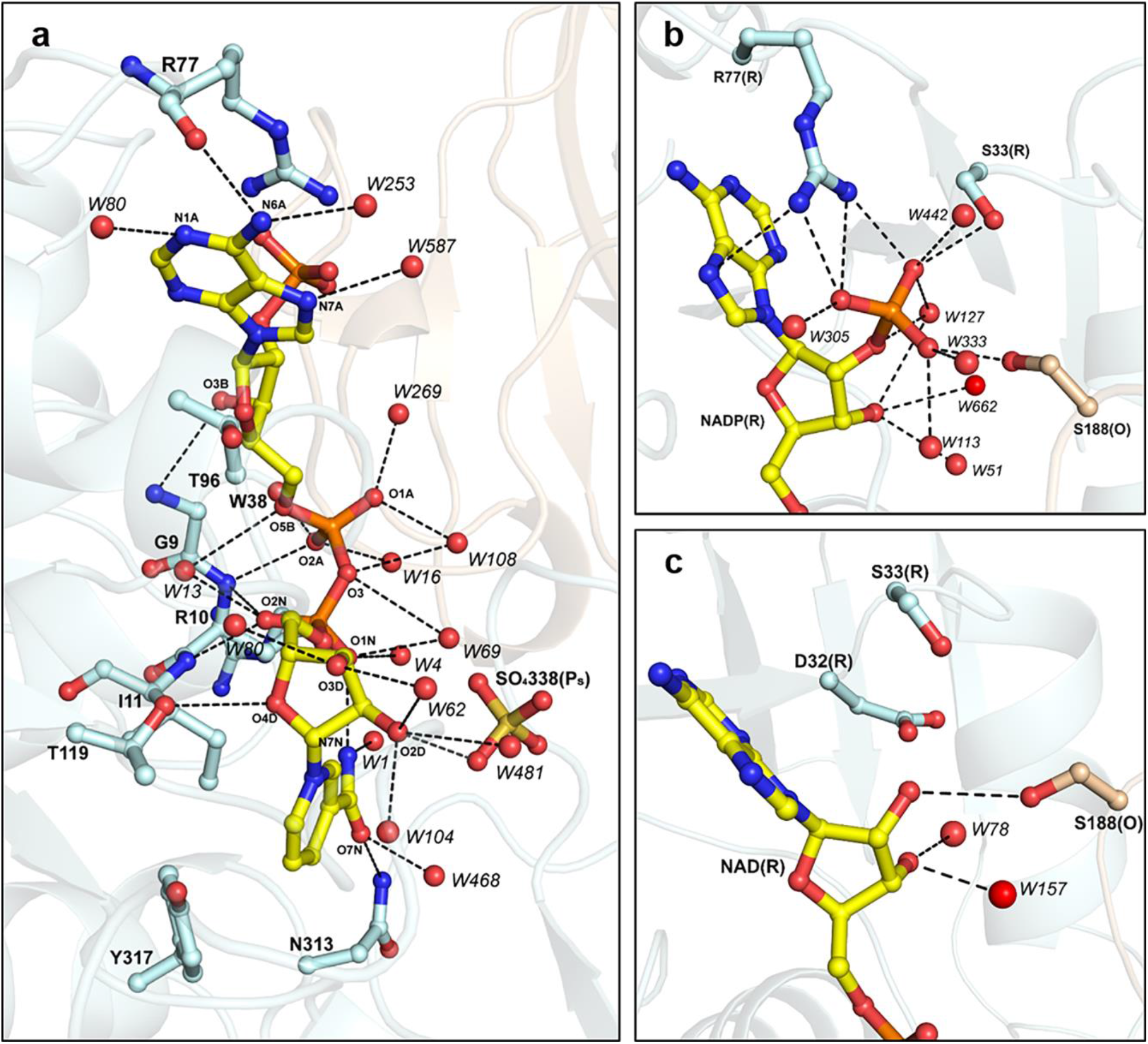
Cofactor interactions in CrGAPA. (**a**) Hydrogen bonds and electrostatic interactions (distance ≤ 3.5 Å) between the CrGAPA cofactor (NADP^+^ or NAD^+^) and protein residues or water molecules. The cofactor bound to chain R of NADP^+^-structure is shown as a representative case. (**b**) Focus on the interactions (distance ≤ 3.5 Å) between the 2’-phosphate group of NADP^+^ and protein residues or water molecules. (**c**) Focus on the interactions (distance ≤ 3.5 Å) between the adenine ribose hydroxyl groups of NAD^+^ and protein residues or water molecules.

The catalytic domain hosts the enzyme active pocket formed by the dyad Cys149/His176, the nicotinamide ring of the cofactor (NADP^+^ or NAD^+^), and two sites named P_s_ and P_i_ hosting the phosphate group(s) of the substrates during catalysis (*i.e.*, BPGA or glyceraldehyde-3-phosphate, G3P) and occupied in all presented structures by sulfate ions coming from the crystallization solution. The reactivity (*i.e.,* nucleophilicity) of the catalytic Cys149 thiol group is ensured by the interaction with the basic imidazole ring of His176 (Cys149 SG – His176 NE2 = 3.3 – 3.4 Å) and by hydrogen bond formation with the backbone amino group and side chain hydroxyl group of Thr150 (Cys149 SG – Thr150 N = 3.2 – 3.3 Å and Cys149 SG – Thr150 OG1 = 3.9 – 4.2 Å) (Figure 3A). The same residues are also responsible of the stabilization of the sulfinylated/sulfonylated Cys149 (Cys-SO_2_^−^/-SO_3_^−^) observed in the oxidized NADP^+^-CrGAPA structure (Supplemental Figure 6).

**Figure 3.**
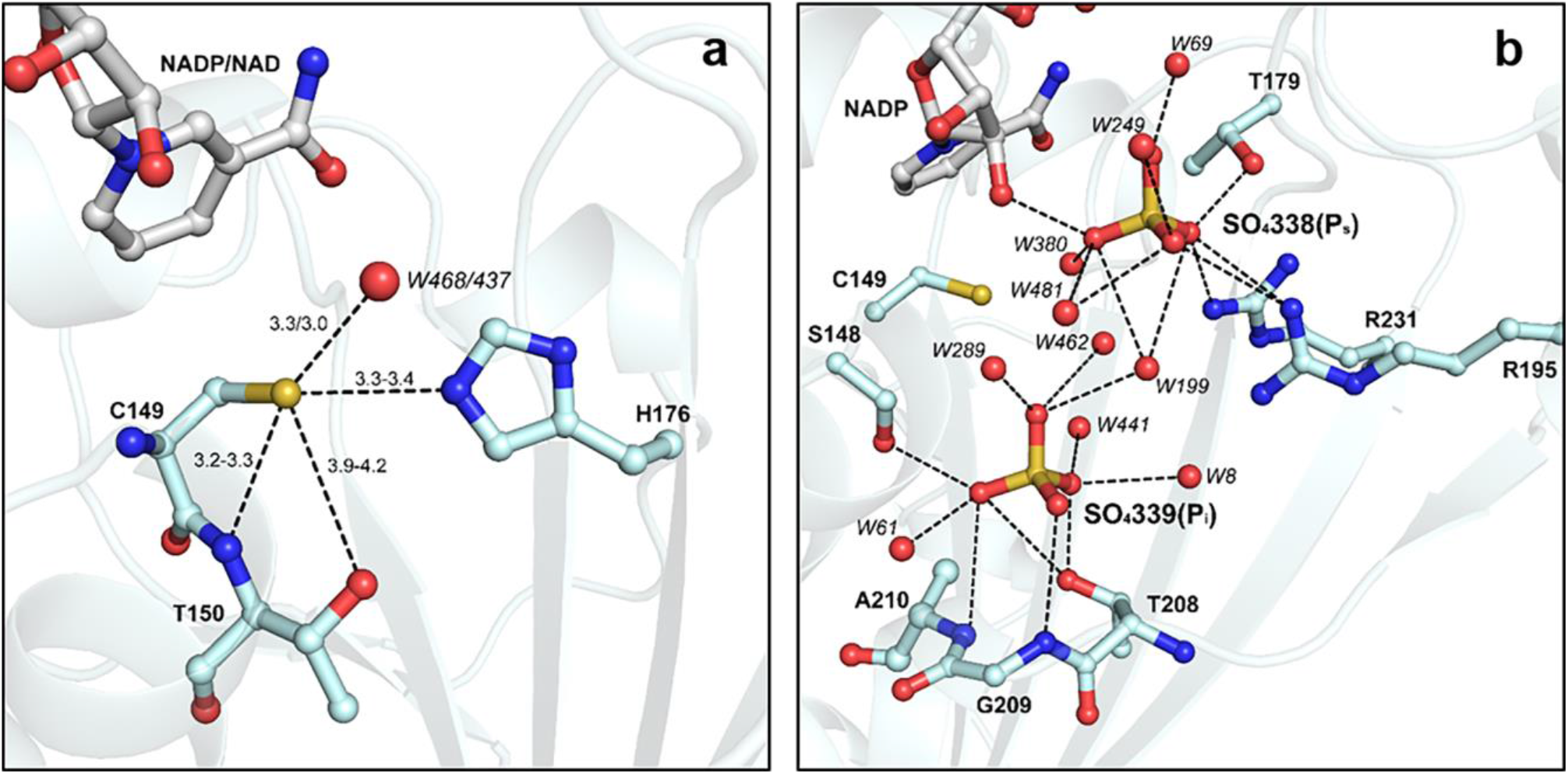
Catalytic site of CrGAPA. (**a**) The catalytic dyad Cys149/His176 and interactions of Cys149 that stabilize the thiolate form, are shown. The ranges of values reported refer to the distances observed in chains O and R of NADP^+^ or NAD^+^ structures. (**b**) The Ps and Pi sites and the hydrogen bonds and electrostatic interactions (distance ≤ 3.5 Å) with protein residues or water molecules, are shown. Ps and Pi sites are occupied in all CrGAPA structures here presented, by sulphate ions coming from the crystallization solution.

The position of the P_S_ and P_I_ sites is superimposable in NADP^+^ and NAD^+^ structures and all residues involved in their stabilization are strictly conserved in photosynthetic GAPDH sequences (Figure 3B and Supplemental Figure 1-3). In particular, the P_S_ site lies close to the cofactor nicotinamide ribose and one of its hydroxyl group interacts with the sulfate ion (Figure 2A). Further stabilization is provided by salt-bridges with Arg195 and Arg231, and by hydrogen bonds with the side chain of Thr179 and water molecules (Figure 3B). The P_I_ site is instead stabilized only by hydrogen bonds with the backbone amino groups of the segment Thr208-Ala210, the hydroxyl groups of Ser148 and Thr208, and water molecules (Figure 3B).

### CrGAPA specifically undergoes S-nitrosylation in the presence of GSNO

In a previous study, we demonstrated that CrGAPA activity is sensitive to oxidative modifications mediated by hydrogen peroxide (H_2_O_2_), GSSG, and GSNO (26). The oxidized NADP^+^-CrGAPA structure here presented, shows that the catalytic Cys is prone to modification even just increasing the pH. Whereas the molecular mechanisms underlying H_2_O_2_- and GSSG-dependent oxidation have been extensively investigated, the nature and type of redox modification induced by GSNO remains elusive.

Here, we analyzed the effect of variable GSNO amounts on CrGAPA activity and we observed a strong inactivation of the enzyme which retains ~30%, 15%, and 10% of residual activity after exposure (30 min) to 0.5, 1, and 2 mM GSNO, respectively (Figure 4A). As mentioned before, CrGAPA has four cysteines showing different accessibility as assessed by their accessible surface area (ASA) (Figure 5). In NADP-CrGAPA, Cys285 is the most exposed (average residue and thiol group ASAs equal to 67 Å^2^ and 20 Å^2^, respectively), while the catalytic Cys149 is less accessible (average residue and thiol group ASAs equal to 9 Å^2^ and 6 Å^2,^ respectively). In contrast, Cys18 and Cys153 are almost buried (average residue and thiol group ASAs equal to 2 Å^2^ and 0 Å^2^ for Cys18 and both 0 Å^2^ for Cys153). Therefore, Cys accessibility values and protein inactivation strongly suggest that GSNO could interact with the catalytic Cys149 and likely Cys285. To establish the specific involvement of the catalytic cysteine, the enzyme was incubated in the presence of the substrate BPGA prior to treatment with GSNO. As shown in Figure 4B, the GSNO-dependent inactivation was almost completely blocked consistently with the fact that BPGA can covalently bind to the catalytic Cys149 and therefore, its presence allows full protection from redox alterations as previously established (23,28). Incubation of CrGAPA with NADP^+^ partially prevented inhibition of the enzyme by GSNO (Figure 4C), suggesting that NADP^+^, bound to the active site, might interfere with the GSNO-dependent inactivation process likely through steric hindrance. Consistently, if the cofactor is removed from the structure the accessibility of Cys149 increases (average residue and thiol group ASAs equal to 28 Å^2^ and 21 Å^2^, respectively).

**Figure 4.**
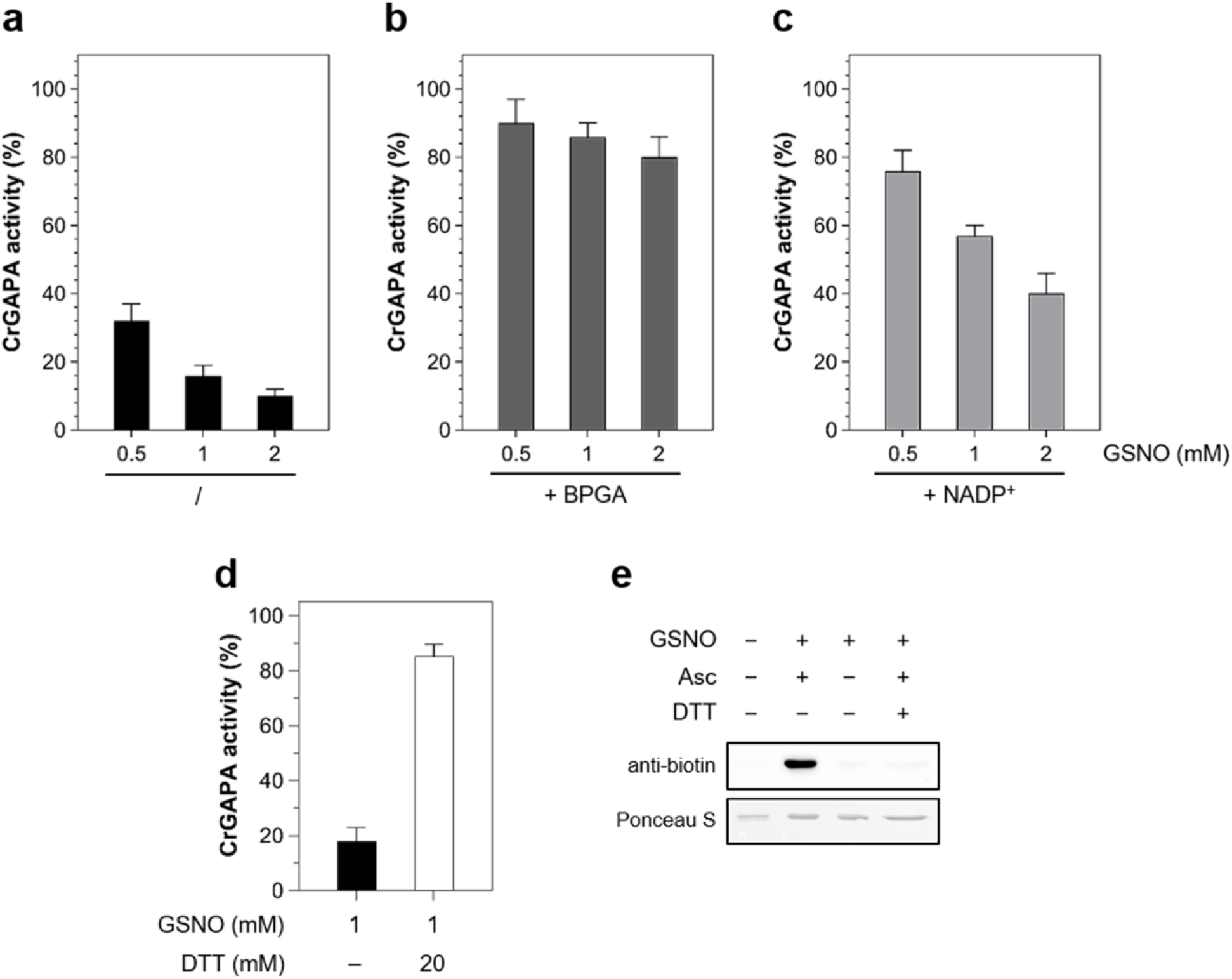
GSNO-mediated S-nitrosylation of CrGAPA. (**a**) Incubation of CrGAPA with GSNO. The enzyme was incubated (30 min) with different concentration of GSNO. Substrate (**b**) and cofactor (**c**) protection of GSNO-treated CrGAPA. CrGAPA was pre-incubated with BPGA-generating system or 0.2 mM NADP^+^ prior to exposure to 2 mM GSNO (see “Material and Methods” fur further details). (**d**) The reversibility of CrGAPA inactivation by GSNO (2 mM, black bar) was assessed after incubation in the presence of 20 mM DTT (white bar). For panels **a-d**, data are represented as mean ± SD (n = 3) of control activity. (**e**) S-nitrosylation of CrGAPA. The enzyme was treated for 30 min in the presence of 2 mM GSNO and nitrosylation was visualized using the BST followed by anti-biotin western blots as described in “Material and Methods”. The red-ponceau (ponceau S) staining of the membrane shows equal loading in each lane.

**Figure 5.**
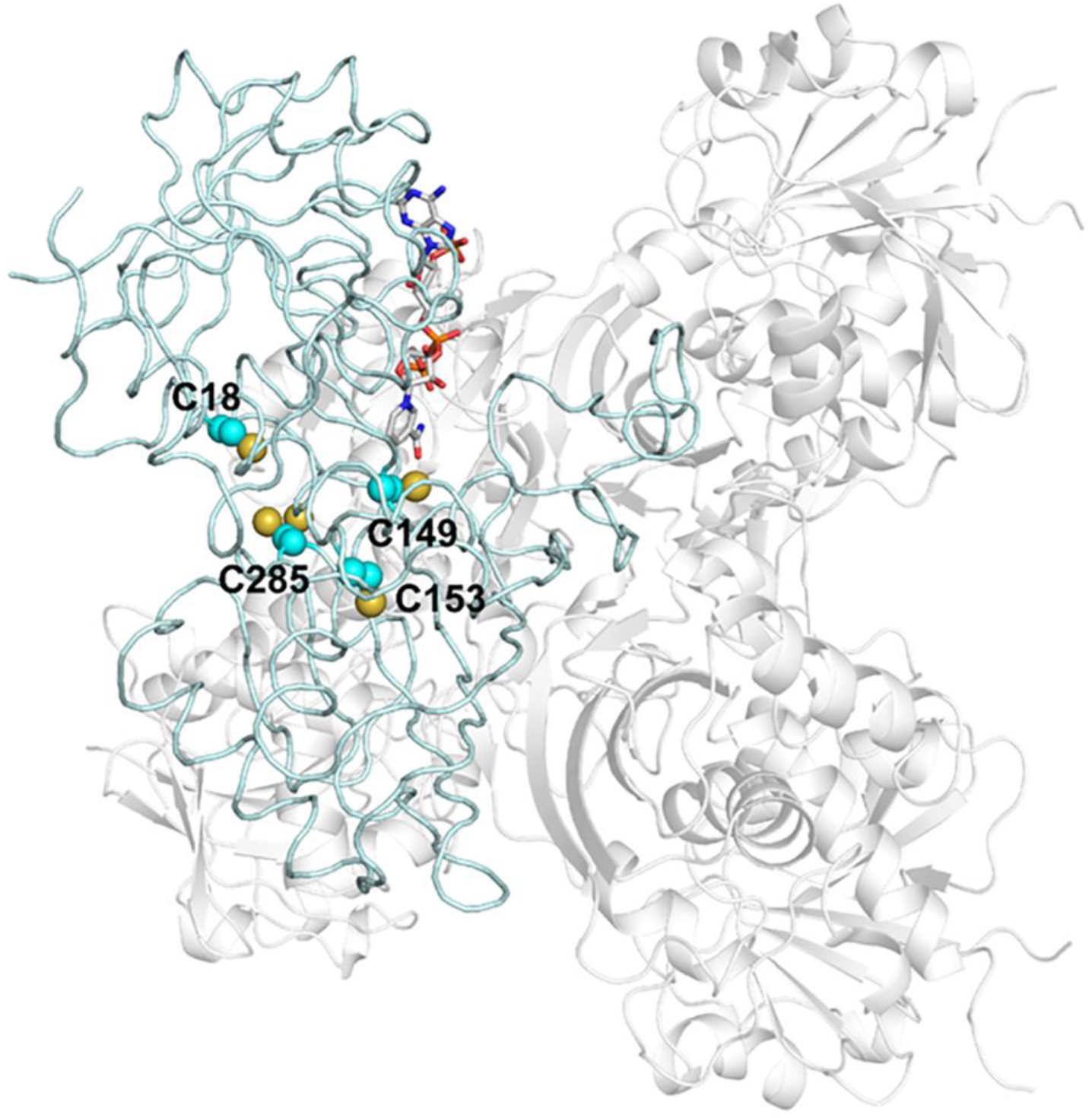
Cysteine residues position and accessibility in CrGAPA monomer. The position of cysteines in the monomer of CrGAPA is shown. Based on accessible surface area (ASA), Cys18 and 153 are buried (2 and 0 Å^2^, and 0 and 0 Å^2^ for average residue and thiol group, respectively), the catalytic Cys149 shows a low accessibility (9 Å^2^ and 6 Å^2^ for average residue and thiol group, respectively), while Cys285 that shows in the crystal structure a double conformation is the most accessible (67 Å^2^ and 20 Å^2^ for average residue and thiol group, respectively).

As aforementioned, GSNO can react with protein thiols to induce S-nitrosylation or S-glutathionylation. Based on activity measurements, we cannot distinguish which of the two redox modifications alter the redox state of CrGAPA catalytic cysteine. To establish the type of redox alteration induced by GSNO, we employed MALDI-TOF mass spectrometry (MS) coupled to the biotin switch technique (BST) and anti-biotin western blots. MALDI-TOF analysis was performed on CrGAPA after incubation with 1 mM GSNO for 30 min (Supplemental Figures 7). Mass spectra for GSNO-treated CrGAPA before or after DTT treatment were identical indicating that GSNO is unable to induce S-glutathionylation. Indeed, S-glutathionylation typically results in a 305 Da shift of the protein mass that can be usually observed by MALDI-TOF MS. By contrast, the labile nitrosothiol is undetectable by MALDI-TOF MS as the NO moiety is lost during the laser-induced ionization process. Consequently, the S-nitrosylated protein is not distinguishable from its unmodified form. Despite the absence of significant mass shift, activity measurements revealed that GSNO causes CrGAPA inhibition (Figure 4E). In addition, reducing treatments almost restored full activity (Figure 4E). Using the biotin-switch technique coupled to anti-biotin western blot, we further evaluated the nitrosylated state of GSNO-treated CrGAPA (Figure 4F). After incubation of CrGAPA for 30 min in the presence of 1 mM GSNO, we observed a strong signal that was absent without ascorbate and completely reversed in the presence of reducing agents (Figure 4F).

Taken together, these results indicate that (i) CrGAPA activity is reversibly inhibited by GSNO; (ii) GSNO causes CrGAPA inhibition solely through S-nitrosylation as revealed by BST and MALDI-TOF MS analyses; (iii) incubation with BPGA fully protects CrGAPA inhibition indicating that the catalytic Cys149 is targeted by GSNO; and (iv) NADP^+^ partially hampers GSNO-dependent S-nitrosylation likely affecting GSNO binding and/or reactivity.

### Structural analysis of GSNO-CrGAPA interactions by molecular dynamics

After establishing that CrGAPA undergoes S-nitrosylation in the presence of GSNO, we carried out molecular dynamics (MD) simulations to gain insight into the dynamics of the GSNO-dependent trans-nitrosylation process of CrGAPA at the molecular level. To evaluate possible variations of the glutathione binding mode prior and after the reaction with the enzyme, we performed two different MD simulations of (i) CrGAPA in complex with GSNO and (ii) S-nitrosylated CrGAPA in complex with glutathione thiolate (GS^-^), *i.e.*, the leaving group formed after the transfer of the NO moiety to the enzyme (4). By a decomposition analysis of the trajectories according to the MM-GBSA scheme, we quantified the contribution of each amino acids to the binding of GSNO/GS^-^, identifying at the atomistic level the GSNO/GS^-^ binding motif.

The binding of GSNO to CrGAPA involves several protein residues including His176, Thr207, Thr208, and Arg231 (Figure 6A). His176 forms the catalytic dyad (Cys149/His176) and it is responsible for the deprotonation of Cys149 and consequent stabilization of the thiolate (-S^-^) (Figure 6B). Therefore, His176 is found in a protonated form with the imidazolium ring bearing two NH groups and a positive net charge. Furthermore, His176 participates to the binding of GSNO as it strongly interacts with the γ-glutamate of GSNO forming a salt bridge and a hydrogen bond with its carboxylate group. Intriguingly, we can note that His176 bridges Cys149 and GSNO, the two groups involved in the NO transfer, by forming two distinct hydrogen bonds through its NH groups. As aforementioned, Thr207, Thr208, and Arg231 also participate in the anchoring process with GSNO interacting with the γ-glutamate group (Figure 6C). Thr207/Thr208 form hydrogen bonds, acting as acceptors, with the N-terminal amino group of GSNO, while Arg231 interacts with the γ-glutamate carboxylate moiety via the typical donor-bifurcated hydrogen bonding/salt bridge. Remarkably, Cys149 by itself has a negative effect on GSNO binding, due to electrostatic repulsion between the negative Cys149 thiolate and glutathione –SNO group.

**Figure 6.**
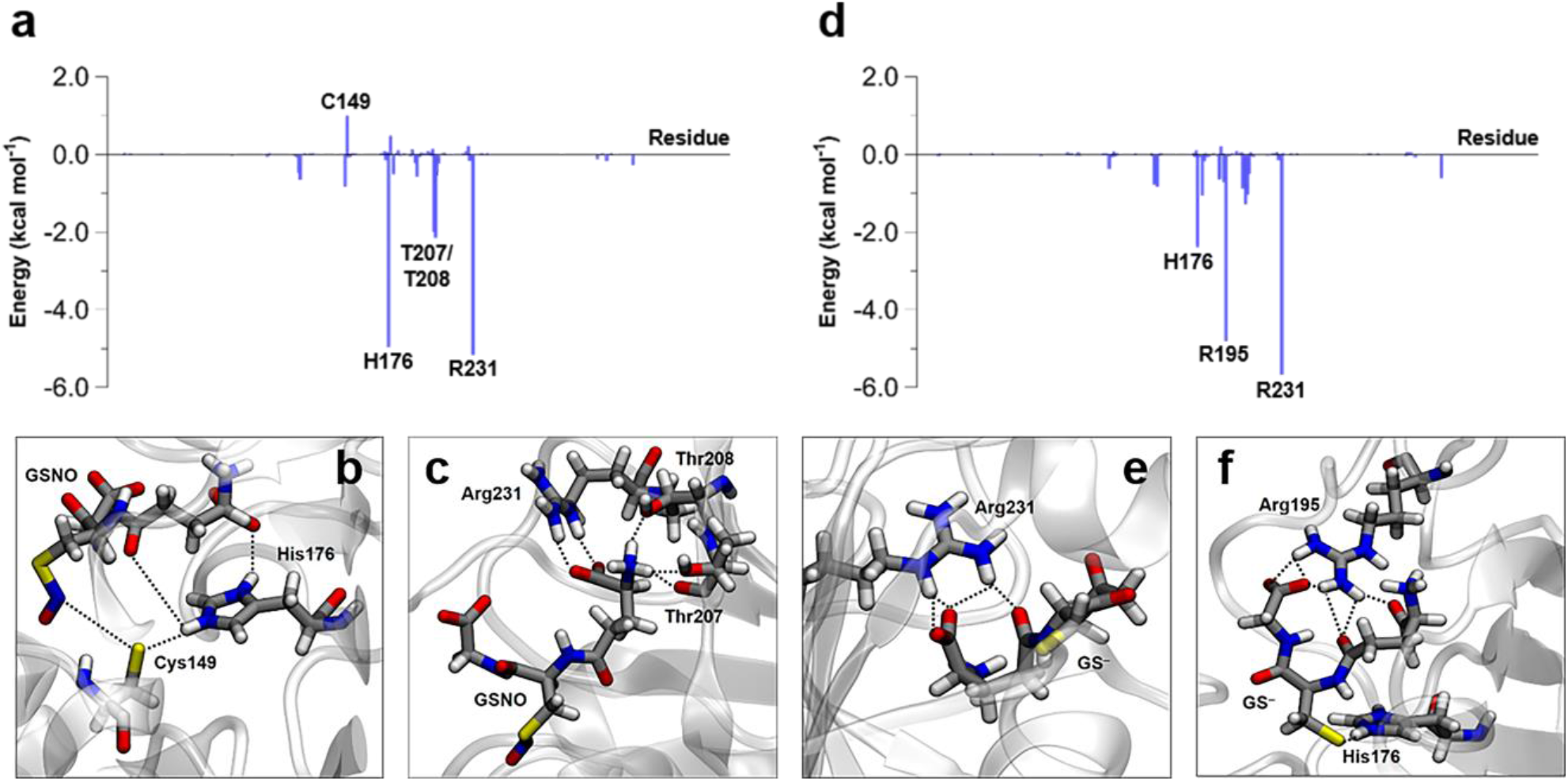
Structural interactions of CrGAPA with GSNO and GS^-^. (**a**) ΔG_binding_ between CrGAPA and GSNO, decomposed per residue. (**b**) Interaction between His176 and GSNO. (**c**) interactions of Thr207, Thr208, Arg231, and GSNO. (**d**) ΔG_binding_ between S-nitrosylated CrGAPA and GS^-^, decomposed per residue. (**e**) Interaction between Arg231 and GS^-^. (**f**) Interactions between His176, Arg195, and GS^-^.

The energy contribution of CrGAPA residues interacting with GS^−^ after the transfer of NO from GSNO to Cys149 has occurred is shown in Figure 6D. Arg231 maintains its interaction with the γ-glutamate of GSNO whereas His176, although still involved GS^−^ binding, undergoes major variations. Notably, Cys149, which originally formed a strong hydrogen bond with His176, loses its negative charge and His176 moves to stabilize by hydrogen bonding the newly formed thiolate in the GS^−^. In the conformational rearrangement induced by the NO transfer, Arg195 intervenes to stabilize the C-terminal carboxylate of the GS^−^ glycine moiety, while Thr207 and Thr208 no longer participate in the stabilization of GS^−^.

As described above, the transfer of the NO moiety from GSNO to Cys149 causes a consistent rearrangement of charges between the two Cys residues involved in the trans-nitrosylation reaction, triggering a reorganization of the network of interactions between the CrGAPA protein and the “GS” scaffold. If GSNO is anchored to CrGAPA mainly by the γ-glutamate residue and kept close to the target Cys149 by His176, a redistribution of the interaction throughout GS^−^ occurs after the trans-nitrosylation reaction. The analysis of the rmsd (*i.e.,* mobility) of the three amino acids comprising the glutathione moiety in the GSNO/CrGAPA(S^−^) and GS^−^/CrGAPA(SNO) complexes, during the MD simulations, neatly reflects this behavior. In the GSNO/CrGAPA complex, γ-glutamate is characterized by a rmsd of 0.76 Å, CysNO of 1.01 Å, and glycine of 1.54 Å, demonstrating that the anchoring of GSNO to the protein mainly involves the γ-glutamate residue. After NO transfer has occurred, the rmsd of glycine decreases to 1.18 Å, while the rmsd values for Cys and γ-glutamate increase (1.23 Å and 1.39 Å, respectively), showing a rigidification of the C-terminal glycine due to the binding with Arg195 and an increase of the flexibility of the Cys and γ-glutamate moieties. Besides, we also noted an increased overall molecular mobility of glutathione which shifts from 1.03 Å for GSNO to 1.29 Å for GS^−^.

### Structural snapshots of the GSNO-dependent S-nitrosylation of CrGAPA

Because MD simulations cannot provide information about reactive processes, a deeper understanding of the trans-nitrosylation reaction was obtained by calculating the energetic profile of the NO transfer from GSNO to Cys149 in the CrGAPA protein environment using a QM/MM approach. Overall, QM/MM calculations are instrumental to get mechanistic insights into reaction profiles providing both thermodynamic parameters (*i.e.*, activation barrier and reaction-free energy) and the relative contribution of protein residues to activation barriers (*i.e.*, fingerprinting analysis). Before analyzing the trans-nitrosylation reaction in the protein microenvironment, we examined the reaction in conventional media (Supplemental Figure 8). While trans-nitrosylation in gas phase is typically a two-step barrierless process (Supplemental Figure 8A), it is a single-step process in water (Supplemental Figure 8B) characterized by an activation energy of 12.6 kcal mol^−1^. Both processes are isoenergetic due to the symmetry of the reagents and products. In contrast, when we considered the protein environment, the transfer of NO from GSNO to Cys149 is an exoergonic single-step process characterized by an activation energy of 3.4 kcal mol^−1^ (Figure 7A). Therefore, the protein environment lowers significantly the barrier of the trans-nitrosylation process and differentiates the energies of the two nitrosylated cysteines, favoring in this case the S-nitrosylation of Cys149. In the transition state (TS), the net negative charge on the sulfur atom of Cys149 is reduced by the approaching of NO and the formation of the incipient S-NO bond, while conversely the cysteine of GS(NO) is becoming a thiolate.

**Figure 7.**
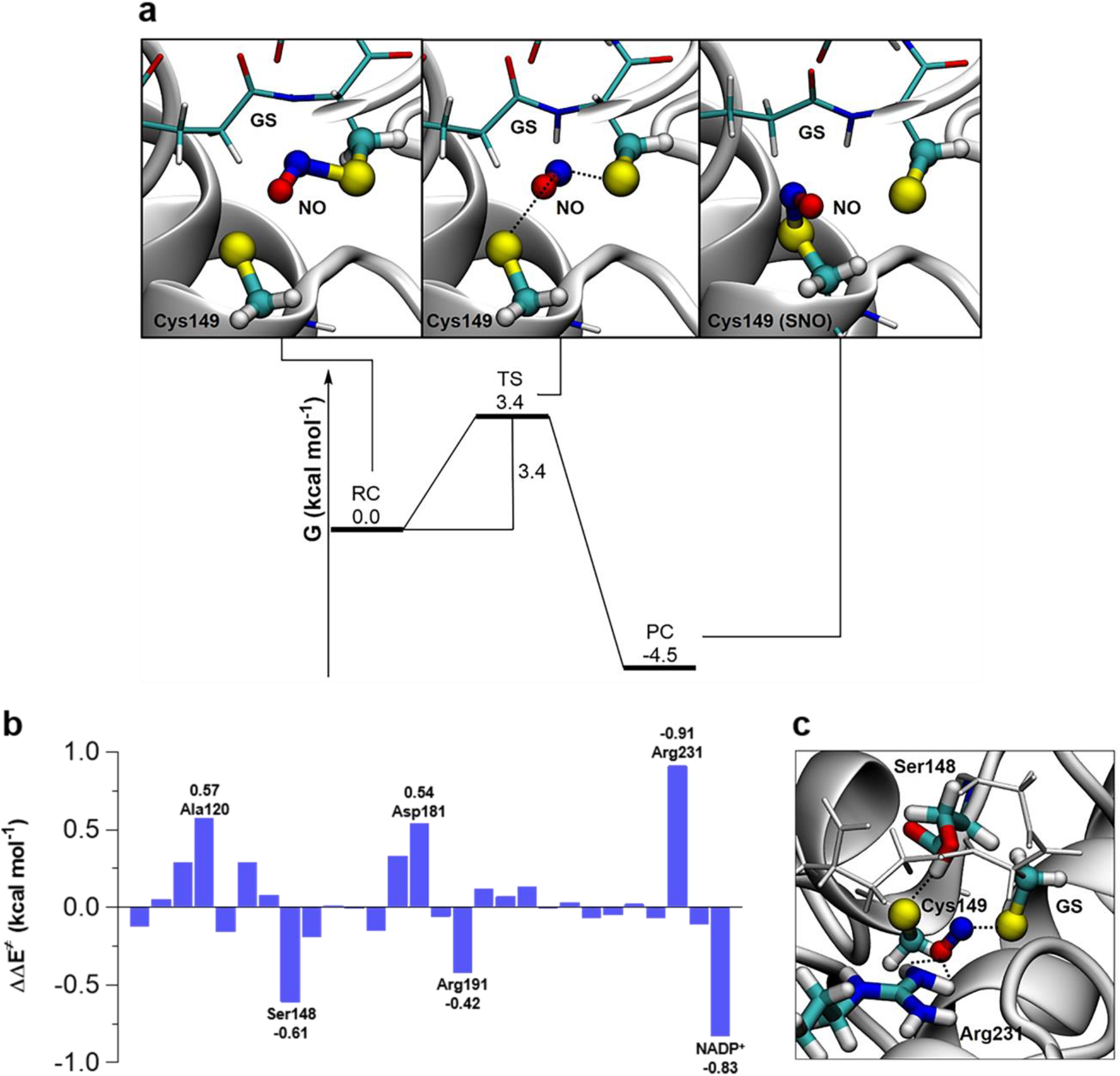
Quantum mechanical analysis of GSNO-dependent S-nitrosylation of CrGAPA. (**a**) Free energy surface (FES) for the trans-nitrosylation process in CrGAPA. Activation barrier and reaction energies are expressed in kcal mol^−1^. In the squares, a blow-up of the critical points. From left to right: reactive complex (RC), transition state (TS), and product complex (PC). (**b**) Stabilizing/destabilizing effect of single amino acids in the activation barrier for the GSNO-dependent trans-nitrosylation process of CrGAPA. (**c**) Interaction between Ser148, Arg231, and the QM reactive system in TS (i.e., Cys149-SNO-GS^−^).

The relative contribution of each specific amino acid to activation barriers was quantified by fingerprint analysis (Figure 7B). Through specific interactions, Ser148 plays an important role in destabilizing the TS while Arg231 has an opposite effect (*i.e.,* stabilization of the TS). Specifically, Arg231 acts as a shuttle that assists the NO moiety moving from GSNO to Cys149 (Figure 7C), stabilizing the TS through hydrogen bonding and electrostatic interactions, as already observed in some transferases (67,68). Before the reaction with GSNO, Ser148 is strongly hydrogen bonded to the thiolate of Cys149. In the TS, however, Cys149 gradually loses its negative charge (Figure 7C) weakening the hydrogen bond with Ser148 resulting in a destabilizing effect on the TS. All the other amino acids identified by the fingerprint analysis are charged residues (Asp181, Arg191) or residues characterized by strong dipoles (C=O and N-H in Ala120), and all are very close to the reaction site. Their net effect is due to the charge rearrangement between the two Cys residues, passing from the reagent to the transition state (activation barrier). In addition to protein residues, also NADP^+^ plays an important role by acting as a destabilizing factor of the TS and thus affecting the S-nitrosylation reaction electrostatically.

## Discussion

Photosynthetic GAPDH is an important enzyme that fulfils major metabolic functions through its participation in the carbon fixation pathway (69). Unlike land plants, Chlamydomonas along with other green microalgae and cyanobacteria only contains one photosynthetic GAPDH isoform composed by identical type A subunits (*i.e.,* GAPA). This isoform differs from photosynthetic AB-GAPDH as it is not subjected to autonomous light-dependent redox regulation. Indeed, GAPA lacks the C-terminal extension found in B subunits that contains the two regulatory cysteines involved in the thioredoxin-dependent dithiol/disulfide interchanges (70,71). As observed for glycolytic isoforms (28), photosynthetic GAPDH can catalyze the enzymatic reaction in both directions and the only catalytic-related difference relies on the specificity towards the cofactors NAD(H) and NADP(H). While glycolytic isoforms exclusively use NAD(H), photosynthetic GAPDH can use both cofactors with a catalytic preference for NADP(H) (22,69).

The enzymatic activity of both photosynthetic and glycolytic GAPDH isoforms is dependent on the presence of a reactive Cys that performs a nucleophilic attack on the substrate. As observed in other GAPDH from plant and non-plant sources, the crystal structure of CrGAPA revealed that the catalytic Cys149 is located in close proximity to a histidine residue (*i.e.,* His176) crucial for thiol deprotonation and thiolate stabilization (Figure 3A). Besides the catalytic Cys149/His176 dyad, CrGAPA shares with other photosynthetic GAPDH an almost identical native folding and a superimposable domain organization. The folding conservation is also accompanied by a strict conservation of residues involved in cofactor(s) and substrate binding. Furthermore, we observed no structural rearrangements related to the accommodation and stabilization of the two cofactors (*i.e.*, NAD^+^ and NADP^+^).

Besides being an important metabolic enzyme, GAPDH is considered a prominent target of thiol switch regulatory mechanisms (22). The reactivity of the catalytic Cys149 makes the protein sensitive to ROS- and RNS-mediated redox modifications, and alteration of the redox state of Cys149 has a direct consequence on enzyme activity. Among redox PTMs, S-nitrosylation plays an important role in providing ubiquitous mechanisms for thiol-mediated regulatory and signaling pathways. This redox modification can indeed modulate protein function and is typically induced by the interaction of reactive Cys thiols with GSNO. Consistently, accumulation of protein nitrosothiols was observed in mutant plants lacking GSNO reductase (GSNOR), the main enzyme controlling the intracellular concentration of GSNO (5,72).

To gain insight into the molecular and atomistic details of S-nitrosylation, we employed photosynthetic CrGAPA which we demonstrated to specifically undergo GSNO-dependent S-nitrosylation on its catalytic cysteine with consequent reversible inhibition of enzyme activity. Based on the here presented NADP^+^-crystal structure, we carried out computational calculations to determine the protein residues contributing to the binding and stabilization of GSNO within the active site. Intriguingly, we observed that the catalytic His176 participates in GSNO accommodation while interacting with the catalytic Cys149 thiolate (Figure 6B). Therefore, His176 appears crucial in NO transfer and in bypassing the electrostatic repulsion between the Cys149 thiolate and the -SNO group of GSNO, both involved in the reaction process. The binding of GSNO also encompasses two threonines (207 and 208) and Arg231, all interacting with the γ-glutamate moiety of GSNO (Figure 6C). A destabilizing interaction between GSNO and Cys149 highlights the crucial role of a GSNO binding site in the protein for the trans-nitrosylation reaction. The chemical recognition of GSNO in this site must provide, through multiple interactions, enough binding energy to the GSNO molecule to bypass the repulsion between the negatively charged catalytic Cys149 and GSNO, a step necessary to activate the trans-nitrosylation process.

The leading role of the γ-glutamate in the positioning and stabilization of the “GS” scaffold was previously observed in the glutathionylated glycolytic AtGAPC1 (31). In the crystal structure of glutathionylated AtGAPC1, the glutathione molecule covalently bound to the catalytic cysteine is set in place by the interactions between several residues that specifically interact with the γ-glutamate moiety, while the C-terminal carboxylate of the glycine moiety is free and seems dispensable in the GSH stabilization.

Here we show that a rearrangement of the interaction network occurs in the binding of glutathione thiolate (GS^−^), which results from the reaction of GSNO with Cys149. This change is functional to the release of GS^−^ and the consequent stabilization of the nitrosothiol on the catalytic Cys149. While Arg231 maintains the bifurcated interaction with the γ-glutamate carboxylate, the two threonines are no longer involved in its stabilization inducing an increase in the mobility of GS^−^ (Figure 6E). Besides, His176 moves away from the neutral nitrosylated Cys149 and interacts with the thiolate of glutathione, while the C-terminal carboxylic group of the glutathionyl glycine interacts with Arg195, which played no role in GSNO binding (Figure 6F). As a result, the leaving group GS^−^ is more mobile than GSNO.

Taken together, these observations highlight the importance of positively charged (His and Arg) and polar residues (Thr) in determining the specificity of CrGAPA S-nitrosylation and their multiple role in this reaction, *i.e.,* deprotonation of the catalytic cysteine, support in NO transfer, and stabilization of the GSNO/GS^−^. Previous studies based on sequence analysis proposed different consensus motifs for S-nitrosylation comprising acidic (*i.e.,* glutamate and aspartate) and basic (*i.e.,* arginine, histidine, and lysine) residues flanking the target cysteine (11,13,73) as well as bioinformatic tools to predict the propensity of a given Cys to undergo S-nitrosylation (74,75). Although predictive, both approaches do not clarify the operative role of the specific residues mainly because they do not consider the three-dimensional structure of the protein. A revised acid-base motif that considers long-distant charged residues has been identified, highlighting the importance of structural and molecular determinants in Cys propensity to S-nitrosylation (12). Despite being proposed in the GSNO binding motif, acidic residues were not detected in the stabilization and accommodation of GSNO in the proximity of CrGAPA Cys149.

QM/MM calculations revealed that the GSNO-dependent trans-nitrosylation of CrGAPA has a ~4-fold lower activation energy compared to the trans-nitrosylation reaction in aqueous solution (Figure 8A and Supplemental Figure 8B). Protein-mediated trans-nitrosylation further differs from the reaction in water as it is an exoergonic process since there is no symmetry between reactants (GSNO + Cys149-S^−^) and products (Cys149-SNO + GS^−^). Thus, the newly formed Cys149-SNO has a lower energy compared to GSNO indicating that the nitrosothiol is more stable in CrGAPA compared to the nitrosylating agent. It is well established that the exothermicity determines the amount of reagent required to induce the reaction, while the height of the energy barrier determines the reaction rate. Therefore, we can hypothesize that, *in vivo*, GSNO (*i.e.*, the reagent) induces a fast CrGAPA S-nitrosylation, regardless of its intracellular concentration.

The activation barrier for the trans-nitrosylation reaction is modulated by the protein microenvironment and fingerprint analysis was instrumental to unravel the relative contribution of CrGAPA residues. Intriguingly, only polar or charged residues (Ser148, Asp181, Arg191, and Arg231) were identified to affect the stability of the transition state (GS…NO…S-Cys149) thus modulating the energy barrier of the reaction (Figure 7B). While Ser148 and Arg191 have a destabilizing effect with the hydroxyl group of Ser148 involved in Cys149 thiolate stabilization, Asp181 and Arg231 contribute to stabilize the TS. Besides protein residues, we found that also NADP^+^ destabilizes the TS and thus increases the energy barrier of the trans-nitrosylation reaction. On this basis, the role of the cofactor in partially preventing the GSNO-dependent CrGAPA inhibition can be ascribed more to electrostatic interactions than to steric hindrance affecting GSNO binding.

In conclusion, we propose a structurally-based computationally-derived GSNO binding motif in which binding and stabilization of GSNO/GS^-^ mainly involve basic and hydroxyl residues that mainly interact with the double charged N-terminal γ-glutamate group (*i.e.*, positive N-terminal amine and negative carboxylic group). The identified residues, namely His176, Arg195, Thr208, and Arg231 (Figure 6), are known to play also an essential role in modulating the catalytic and redox reactivity of Cys149 and in the stabilization of the substrate. Thus, it appears clear that catalytic properties along with redox sensitivity to GSNO-mediated oxidation are controlled/operated by the same network of residues. Moreover, we found that the energy profile of CrGAPA S-nitrosylation is modulated by the native protein environment involving both short- and long-distance electrostatic and polar interactions (Figure 7). The importance of charge interconnections, even at long distance, was previously observed in Arabidopsis GAPC1 and GAPA, where they contributed to tune cysteine reactivity towards H_2_O_2_-dependent primary oxidation (*i.e.,* sulfenic acid formation) (27).

Albeit cysteines are unique molecular switches and highly responsive sensors of the cellular redox state, the molecular mechanisms underlying thiol oxidative modifications are not fully elucidated. Computational analyses coupled to structural and biochemical studies appear essential to the understanding of the oxidation sensitivity of reactive Cys and the complex mechanisms underlying oxidative modifications (74), which are fundamental PTMs for the functioning and regulation of cellular networks alongside other more studied modifications such as phosphorylation.

Here we demonstrated that GSNO-mediated nitrosothiol formation affects the functioning of photosynthetic CrGAPA. However, the physiological impact of S-nitrosylation on the algal enzyme remains to be investigated and related to this, also the NO-dependent redox modulation of the carbon fixation pathway in microalgae and other photosynthetic organisms is still unexplored. To note, all enzymes participating in the carbon fixation pathway were identified as putative targets of S-nitrosylation (19) but molecular evidence for NO-dependent regulation of CBB enzymes is still limited (26,76). To date, only GAPA was demonstrated to undergo S-nitrosylation acting through a thiol-based regulatory mechanism of protein activity. The sensitivity of GAPDH to multiple types of redox PTMs might be exploited to measure precisely the occurrence, extent, and reversibility of a given redox modifications. In addition, we should mention that moonlighting functions of animal GAPDH are specifically triggered by redox modifications of the catalytic cysteine which redirect the enzyme to new and completely unrelated functions (22). In particular, nuclear translocation of nitrosylated GAPDH can control apoptosis but also regulation of gene expression and it was demonstrated to act as a trans-nitrosylase of nuclear proteins (22). Whether photosynthetic GAPDH also possesses additional functions is still an open question and further studies are required to shed light on the possibility that this enzyme might be involved in S-nitrosylation-dependent regulatory cascades in green algae and other photosynthetic organisms.

## Acknowledgments

We gratefully acknowledge Elettra (Trieste) and the staff of XRD1 beam line for allocation of beam time. SF and GF thank the Consorzio Interuniversitario di Ricerca in Chimica dei Metalli nei Sistemi Biologici (CIRCMSB). This work was supported by University of Bologna Alma Idea 2017 Program (to MZ);CNRS Sorbonne Université, Agence Nationale de la Recherche Grant 17-CE05-0001 CalvinDesign (to CHM and SDL); LABEX DYNAMO (ANRLABX-011 to CHM, MDM, and SDL) and EQUIPEX CACSICE (ANR-11-EQPX-0008 to CHM and SDL), partly through funding of the Proteomic Platform IBPC (PPI). JR, MM and SFanti are supported by a PhD grant from the University of Bologna (PhD programs in Cellular and Molecular Biology for JR and MM and in Nanoscience for Medicine and the Environment for SFanti).

## Author contribution

MC, SFermani, and MZ designed the research; EJM, JR, MM, SFanti, CHM, MDM, SFermani, and MZ performed the research; EJM, JR, MM, MC, SFermani, and MZ analyzed the data; and EJM, JR, MC, SFermani, and MZ wrote the paper. All authors have read and agreed to the current version of the manuscript.

