## Supplemental Material for "Structural snapshots of nitrosoglutathione binding and reactivity underlying S-nitrosylation of photosynthetic GAPDH"

**SUPPORTING INFORMATION**

Supporting information of this manuscript includes eight figures and one table:

**Figure S1.** Sequence alignment of CrGAPA and photosynthetic GAPA from land plants.

**Figure S2.** Sequence alignment of CrGAPA and photosynthetic GAPA from green algae.

**Figure S3.** Sequence alignment of CrGAPA and photosynthetic GAPA from cyanobacteria.

**Figure S4.** SDS-PAGE of CrGAPA.

**Figure S5.** Interactions of His-tag within $NADP^+$-CrGAPA crystal structure

**Figure S6.** Interactions of the catalytic sulfnylated/sulfonylated Cys149 within oxidized $NADP^+$-CrGAPA crystal structure

**Figure S7.** MALDI-TOF mass spectrometry analysis of CrGAPA

**Figure S8.** Free energy surface (FES) for the trans-nitrosylation process

**Table S1.** Refinement statistics for CrGAPA crystal structures

### Figure S1

```
                   1     8         18                32
CrGAPA      EKKIRVAINGFGRIGRNFLRCWHGRQN-TLLDVVAINDSGGVKQASHLLKYDSTLGTFAA
AtGAPA1     --KLKVAINGFGRIGRNFLRCWHGRKD-SPLDIIAINDTGGVKQASHLLKYDSTLGIFDA
AtGAPA2     --KIKVAINGFGRIGRNFLRCWHGRKD-SPLDVVVINDTGGVKQASHLLKYDSTLGIFDA
SoGAPA      --KLKVAINGFGRIGRNFLRCWHGRKD-SPLDVVVINDTGGVKQASHLLKYDSILGTFDA
NtGAPA      --KLKVAINGFGRIGRNFLRCWHGRKD-SPLDVIAINDTGGVKQASHLLKYDSTLGIFDA
OsGAPA      --KLKVAINGFGRIGRNFLRCWHGRGDSSPLDVIAINDTGGVKQASHLLKYDSTLGIFDA
StGAPA      --KLKVAINGFGRIGRNFLRCWHGRKD-SPLDVIAINDTGGVKQASHLLKYDSTLGIFDA
MtGAPA      --KIKVAINGFGRIGRNFLRCWHGRKD-SPLDVIAINDTGGVKQASHLLKYDSTLGIFDA
ZmGAPA      --KLKVAINGFGRIGRNFLRCWHGRGDASPLDVIAINDTGGVKQASHLLKYDSTLGIFDA
              *::******************* : : **::.***:************** ** * *

                                      77                 96
CrGAPA      DVKIVDDSHISVDGKQIKIVSSRDPLQLPWKEMNIDLVIEGTGVFIDKVGAGKHIQAGAS
AtGAPA1     DVKPSGETAISVDGKIIQVVSNRNPSLLPWKELGIDIVIEGTGVFVDREGAGKHIEAGAK
AtGAPA2     DVKPSGDSALSVDGKIIKIVSDRNPSNLPWGELGIDLVIEGTGVFVDRDGAGKHLQAGAK
SoGAPA      DVKTAGDSAISVDGKVIKVVSDRNPVNLPWGDMGIDLVIEGTGVFVDRDGAGKHLQAGAK
NtGAPA      DVKPVGTDGISVDGKVIQVVSDRNPVNLPWGDLGIDLVIEGTGVFVDREGAGKHIQAGAK
OsGAPA      DVKPVGDNAISVDGKVIKVVSDRNPSNLPWGELGIDLVIEGTGVFVDRDGAGKHIQAGAK
StGAPA      DVKPVGTDGISVDGKIIQVVSNRDPVNLPWGELGIDLVIEGTGVFVDREGAGKHIQAGAK
MtGAPA      DVKPVGTDGISVDGKVIKVVSDRNPANLPWGELGIDLVIEGTGVFVDREGAGKHITAGAK
ZmGAPA      DVKPVGDNAISVDGKVIKVVSDRNPSNLPWGELGIDLVIEGTGVFVDREGAGKHIQAGAK
            ***    .    :***** *::**.*:*   ***  ::.**:********:*:  *****: ***.

              119                                 149 153
CrGAPA      KVLITAPAKDKDIPTFVVGVNEGDYKHEYPIISNASCTTNCLAPFVKVLEQKFGIVKGTM
AtGAPA1     KVIITAPGK-GDIPTYVVGVNADAYSHDEPIISNASCTTNCLAPFVKVLDQKFGIIKGTM
AtGAPA2     KVLITAPGK-GDIPTYVVGVNAELYSHEDTIISNASCTTNCLAPFVKVLDQKFGIIKGTM
SoGAPA      KVLITAPGK-GDIPTYVVGVNEEGYTHADTIISNASCTTNCLAPFVKVLDQKFGIIKGTM
NtGAPA      KVLITAPGK-GDIPTYVVGVNADLYNPDEPIISNASCTTNCLAPFVKVLDQKFGIIKGTM
OsGAPA      KVLITAPGK-GDIPTYVVGVNADQYSPDEPIISNASCTTNCLAPFVKILDQKFGIIKGTM
StGAPA      KVLITAPGK-GDIPTYVVGVNAELYNHDEPIISNASCTTNCLAPFVKVLDQKFGIIKGTM
MtGAPA      KVLITAPGK-GDIPTYVVGVNADGYSHADDIISNASCTTNCLAPFVKVLDQKFGIIKGTM
ZmGAPA      KVLITAPGK-GDIPTYVVGVNADQYNPDEPIISNASCTTNCLAPFVKVLDQKFGIIKGTM
            **:****.*  ****:*****    *.    ****************:*:*****:****

             176         188   195          208                   231
CrGAPA      TTTHSYTGDQRLLDASHRDLRRARAAALNIVPTTTGAAKAVSLVLPSLKGKLNGIALRVP
AtGAPA1     TTTHSYTGDQRLLDASHRDLRRARAAALNIVPTSTGAAKAVALVLPNLKGKLNGIALRVP
AtGAPA2     TTTHSYTGDQRLLDASHRDLRRARAAALNIVPTSTGAAKAVALVLPNLKGKLNGIALRVP
SoGAPA      TTTHSYTGDQRLLDASHRDLRRARAACLNIVPTSTGAAKAVALVLPNLKGKLNGIALRVP
NtGAPA      TTTHSYTGDQRLLDASHRDLRRARAAALNIVPTSTGAAKAVALSSQALRGSSMALPLRVP
OsGAPA      TTTHSYTGDQRLLDASHRDLRRARAAALNIVPTSTGAAKAVALVLPNLKGKLNGIALRVP
StGAPA      TTTHSYTGDQRLLDASHRDLRRARAAALNIVPTSTGAAKAVALVLPSLKGKLNGIALRVP
MtGAPA      TTTHSYTGDQRLLDASHRDLRRARAAALNIVPTSTGAAKAVALVLPSLKGKLNGIALRVP
ZmGAPA      TTTHSYTGDQRLLDASHRDLRRARAAALNIVPTSTGAAKAVSLVLPNLKGKLNGIALRVP
            *************************.******:*******:*    *:*.  .: ****

                                                                 285
CrGAPA      TPTVSVVDLVVQVEKKTFAEEVNAAFREAANGPMKGVLHVEDAPLVSIDFKCTDQSTSID
AtGAPA1     TPNVSVVDLVVQVSKKTFAEEVNAAFRDSAEKELKGILDVCDEPLVSVDFRCSDFSTTID
AtGAPA2     TPNVSVVDLVVQVSKKTFAEEVNAAFRDAAEKELKGILDVCDEPLVSVDFRCSDVSSTID
SoGAPA      TPNVSVVDLVVQVSKKTFAEEVNAAFRESADNELKGILSVCDEPLVSIDFRCTDVSSTID
NtGAPA      TPNVSVVDLVVQVSKKTFAEEVNAAFREAADKELKGILDVCDEPLVSVDFRCSDVSSTVD
OsGAPA      TPNVSVVDLVVQVSKKTLAEEVNQAFRDSAANELKGILEVCDVPLVSVDFRCSDVSCTID
StGAPA      TPNVSVVDLVVQVTKKTFAEEVNAAFREAADKELNGILAVCDEPLVSVDFRCSDVSSTID
MtGAPA      TPNVSVVDLVVQVSKKTFAEEVNEAFRDSAAKELSGILSVCDEPLVSVDFRCTDVSSTVD
ZmGAPA      TPNVSVVDLVVQVSKKTLAEEVNQAFRDAAANELTGILEVCDVPLVSVDFRCSDVSSTID
            **.********** ***:***** ***::*    :.*:* * * ****:**:*:* * ::*

                              313 317
CrGAPA      ASLTMVMGDDMVKVVAWYDNEWGYSQRVVDLAEVTAKKWVA
AtGAPA1     SSLTMVMGDDMVKVIAWYDNEWGYSQRVVDLADIVANNWK-
AtGAPA2     SSLTMVMGDDMVKVIAWYDNEWGYSQRVVDLADIVANNWK-
SoGAPA      SSLTMVMGDDMVKVIAWYDNEWGYSQRVVDLADIVANKWQ-
NtGAPA      ASLTMVMGDDMVKVIAWYDNEWGYSQRVVDLADIVANQWK-
OsGAPA      ASLSMVMGDDMVKVIAWYDNEWGYSQRVVDLADIVANQWK-
StGAPA      SSLTMVMGDDMVKVIAWYDNEWGYSQRVVDLADIVANQWK-
MtGAPA      SSLTMVMGDDMVKVIAWYDNEWGYSQRVVDLADIVANNWK-
ZmGAPA      ASLTMVMGDDMVKVISWYDNEWGYSQRVVDLADICANQWK-
            :**:**********::****************::  *::*
```

**Figure S1. Sequence alignment of CrGAPA and photosynthetic GAPA from land plants.** Abbreviation and UniProtKB entries: CrGAPA, *Chlamydomonas reinhardtii* (Cr) GAPA, A8HP84; AtGAPA1, *Arabidopsis thaliana* (At) GAPA1, P25856; AtGAPA2, *Arabidopsis thaliana* (At) GAPA2, Q9LPW0; SoGAPA, *Spinacia oleracea* (So) GAPA, P19866; NtGAPA, *Nicotiana tabacum* (Nt) GAPA, P09043; OsGAPA, *Oryza sativa* (Os) GAPA, Q7X8A1; StGAPA, *Solanum tuberosum* (St) GAPA, M1ATQ7; MtGAPA, *Medicago truncatula* (Mt) GAPA, G7KYC0; ZmGAPA, *Zea mays* (Zm) GAPA, A0A1D6E965. Residues numbering is according to the crystal structure of CrGAPA. The catalytic dyad is indicated on a black background. Residues involved in cofactor and substrate binding are indicated on a red and blue background. Cysteine residues with the exception of Cys149 are indicated on a light grey background. Invariant residues are marked by an asterisk conserved residues while single or double point indicates conservation between residues with weakly or strongly similar properties, respectively. The proteins were aligned using Clustal Omega (http://www.ebi.ac.uk/Tools/msa/clustalo/).

### Figure S2

```
               1      8        18              32
CrGAPA      EKKIRVAINGFGRIGRNFLRCWHGRQNTLLDVVAINDSGGVKQASHLLKYDSTLGTFAAD
VcGAPA      EKKIRVAINGFGRIGRNFLRCWHGRQNTLLDVVAINDSGGVKQASHLLKYDSTLGTFDAD
OtGAPA      GKTIKVAINGFGRIGRNFLRCWHGRANTMLDIVAINDSGGVKQASHLVKYDSVLGTFEAD
MpGAPA      KAKIKVAINGFGRIGRNFLRCWHGRKDSLLDVVAINDSGGVKQASHLVKYDSILGTFQAD
              .*:******************** :::**:***************:**** **** **

                                77                 96
CrGAPA      VKIVDDSHISVDGKQIKIVSSRDPLQLPWKEMNIDLVIEGTGVFIDKVGAGKHIQAGASK
VcGAPA      VKIVDDSHISVDGKVIKIVSSRDPLQLPWGEMDVDLVIEGTGVFIDKVGAGKHIQAGAKK
OtGAPA      VKIVDDTHISIDGKKIEIVSSRDPLQLPWKALGVDLVIEGTGVFIDTPGASKHITAGASK
MpGAPA      VKIVDDTHMSIDGKSIQIVSSRDPLQLPWKELGVQIVIEGTGVFIDTPGASKHLTAGASK
            ******:*:*:*** *:************  :.:::**********. **.**: ***.*

             119                             149 153
CrGAPA      VLITAPAKDKDIPTFVVGVNEGDYKHEYPIISNASCTTNCLAPFVKVLEQKFGIVKGTMT
VcGAPA      VLITAPAKDKDIPTFVVGVNEADYKHEYPIISNASCTTNCLAPFVKVLQEKFGIVKGTMT
OtGAPA      VLITAPAKGDDIPTYVLGVNADGYKHSDKIISNASCTTNGLAPFVKVIDEKFGLVKGLMT
MpGAPA      VIITAPAKGDDIPTYVMGVNADDYKDSDKIISNASCTTNGLAPFAKVLDDSFGIIKGLMT
            *:******..****:*:***  .**..  ********** ****.**:::.**::** **

            176         188   195          208                    231
CrGAPA      TTHSYTGDQRLLDASHRDLRRARAAALNIVPTTTGAAKAVSLVLPSLKGKLNGIALRVPT
VcGAPA      TTHSYTGDQRLLDASHRDLRRARAAALNIVPTTTGAAKAVALVLPQLKGKLNGIALRVPT
OtGAPA      TTHSYTGDQRILDASHRDLRRARAACLNIVPTSTGAAKAVALVLPQLKGKLNGIALRVPT
MpGAPA      TTHSYTGDQRILDASHRDLRRARAAALNIVPTSTGAAKAVALVLPQLKGKLNGIALRVPT
            **********:**************.******:*******:****.**************

                                                            285
CrGAPA      PTVSVVDLVVQVEKKTFAEEVNAAFREAANGPMKGVLHVEDAPLVSIDFKCTDQSTSIDA
VcGAPA      PTVSVVDLVVQVEKKTFAEEVNAAFREAAAGPMKGVLHVEDAPLVSIDFKCTDQSTSIDS
OtGAPA      PNVSVVDLVVQTSKKCTADEVNAAFREAANGPLKGILAVADEPLVSCDFRCTDVSCTIDA
MpGAPA      PNVSVVDMVIQVEKKCTAEDVNAAFREAAAGPMKGILQVVDEPLVSIDFRCTDVSTSIDS
            *.*****:*:*..**  *::********* **:**:* * * **** **:*** * :**:

                            313 317
CrGAPA      SLTMVMGDDMVKVVAWYDNEWGYSQRVVDLAEVTAKKWVA-
VcGAPA      SLTMVMGDDMVKVVAWYDNEWGYSQRVVDLAEVTAQKWVA-
OtGAPA      ALTMVMGDDMLKVVSWYDNEWGYSQRVVDLAELCAANWE--
MpGAPA      ALTMVMGEDLVKIVAWYDNEWGYSQRVVDLAELTASKWGSK
            :******:*::*:*:*****************: * :*
```

**Figure S2. Sequence alignment of CrGAPA and photosynthetic GAPA from green algae.** Abbreviation and UniProtKB entries: CrGAPA, *Chlamydomonas reinhardtii* (Cr) GAPA, A8HP84. VcGAPA, *Volvox carteri f. nagariensis* (Vc) GAPA, D8U9J4; OtGAPA, *Ostreococcus tauri* (Ot) GAPA, A5H1G5; MpGAPA, *Micromonas pusilla* (Mp) GAPA, C1MIF7. Residues numbering is according to the crystal structure of CrGAPA. The catalytic dyad is indicated on a black background. Residues involved in cofactor and substrate binding are indicated on a red and blue background. Cysteine residues with the exception of Cys149 are indicated on a light grey background. Invariant residues are marked by an asterisk conserved residues while single or double point indicates conservation between residues with weakly or strongly similar properties, respectively. The proteins were aligned using Clustal Omega (http://www.ebi.ac.uk/Tools/msa/clustalo/).

### Figure S3

```
                1      8        18                32
CrGAPA      EKKIRVAINGFGRIGRNFLRCWHGRQ-NTLLDVVAINDSGGVKQASHLLKYDSTLGTFAA
SeGAPA      -MTIRVAINGFGRIGRNFLRCWFGRQ-NTDLEVVAINNTSDARTAAHLLEYDSVLGRFN-
TeGAPA      --MVRVAINGFGRIGRNFMRCWLQRKANSKLEIVGINDTSDPRTNAHLLKYDSMLGIFQD
               :**************:***  *: *: *::*.**::.. :  :***:*** ** *

                                 77                  96
CrGAPA      DVKIVDDSHISVDGKQIKIVSSRDPLQLPWKEMNIDLVIEGTGVFIDKVGAGKHIQAGAS
SeGAPA      ADISYDENSITVNGKTMKIVCDRNPLNLPWKEWDIDLVIESTGVFVTAEGASKHIQAGAK
TeGAPA      AEITADDDCIYAGGHAVKCVSDRNPENLPWSAWGIDLVIEATGVFTSREGASKHLSAGAK
                 *:. * ..*: :* *..*:* :***.  .******.****    **.**:.***.

               119                             149 153
CrGAPA      KVLITAPAKDKDIPTFVVGVNEGDYKHE-YPIISNASCTTNCLAPFVKVLEQKFGIVKGT
SeGAPA      KVLITAPGKAEGVGTYVIGVNDSEYRHEDFAVISNASCTTNCLAPVAKVLHDNFGIIKGT
TeGAPA      KVLITAPGKG-NIPTYVVGVNHHTYDPSE-DIVSNASCTTNCLAPIVKVLHEAFGIQQGM
            *******.*  .: *:*:***.  *  .   ::************..***.: *** :*

              176         188   195          208                  231
CrGAPA      MTTTHSYTGDQRLLDASHRDLRRARAAALNIVPTTTGAAKAVSLVLPSLKGKLNGIALRV
SeGAPA      MTTTHSYTLDQRILDASHRDLRRARAAAVNIVPTTTGAAKAVALVIPELKGKLNGIALRV
TeGAPA      MTTTHSYTGDQRLLDASHRDLRRARAAAMNIVPTSTGAAKAVGLVIPELQGKLNGIALRV
            ******** ***:***************:*****:*******.**:*.*:**********

                                                               285
CrGAPA      PTPTVSVVDLVVQVEKKTFAEEVNAAFREAANGPMKGVLHVEDAPLVSIDFKCTDQSTSI
SeGAPA      PTPNVSVVDLVVQVEKPTITEQVNEVLQKASQTTMKGIIKYSDLPLVSSDFRGTDESSIV
TeGAPA      PTPNVSVVDFVAQVEKPTIAEQVNQVIKEASETTMKGIIHYSELELVSSDYRGHNASSIL
            ***.*****:*.**** *::*:** .:::*::  ***::: .:  *** *::  : *: :

                              313 317
CrGAPA      DASLTMVMGDDMVKVVAWYDNEWGYSQRVVDLAEVTAKKWVA------------------
SeGAPA      DSSLTLVMDGDLVKVIAWYDNEWGYSQRVVDLAELAARKSGRLGLVQTLLISSQDSLPLT
TeGAPA      DASLTMVLGGNLVKVVAWYDNEWGYSQRVLDLAEHMAAHWA-------------------
            *:***:*:..::***:*************:****  * :

CrGAPA      -----------------------
SeGAPA      GGFFVAKREHPRSQDSINPSGPD
TeGAPA      -----------------------
```

**Figure S3. Sequence alignment of CrGAPA and photosynthetic GAPA from cyanobacteria.** Abbreviation and UniProtKB entries: CrGAPA, *Chlamydomonas reinhardtii* (Cr) GAPA, A8HP84. SeGAPA, *Synechococcus elongatus* (Se) GAPA, Q9R6W2; TeGAPA, *Thermosynechococcus elongatus* (Te) GAPA, Q8DMR8. Residues numbering is according to the crystal structure of CrGAPA. The catalytic dyad is indicated on a black background. Residues involved in cofactor and substrate binding are indicated on a red and blue background. Cysteine residues with the exception of Cys149 are indicated on a light grey background. Invariant residues are marked by an asterisk conserved residues while single or double point indicates conservation between residues with weakly or strongly similar properties, respectively. The proteins were aligned using Clustal Omega (http://www.ebi.ac.uk/Tools/msa/clustalo/).

### Figure S4

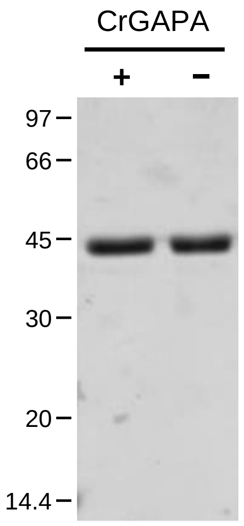


**Figure S5. SDS-PAGE of recombinant CrGAPA**. Sample proteins (3 µg) were separated by 12% polyacrylamide gel under reducing (+) or non-reducing (−) conditions and stained with Coomassie Brilliant Blue.

### Figure S5

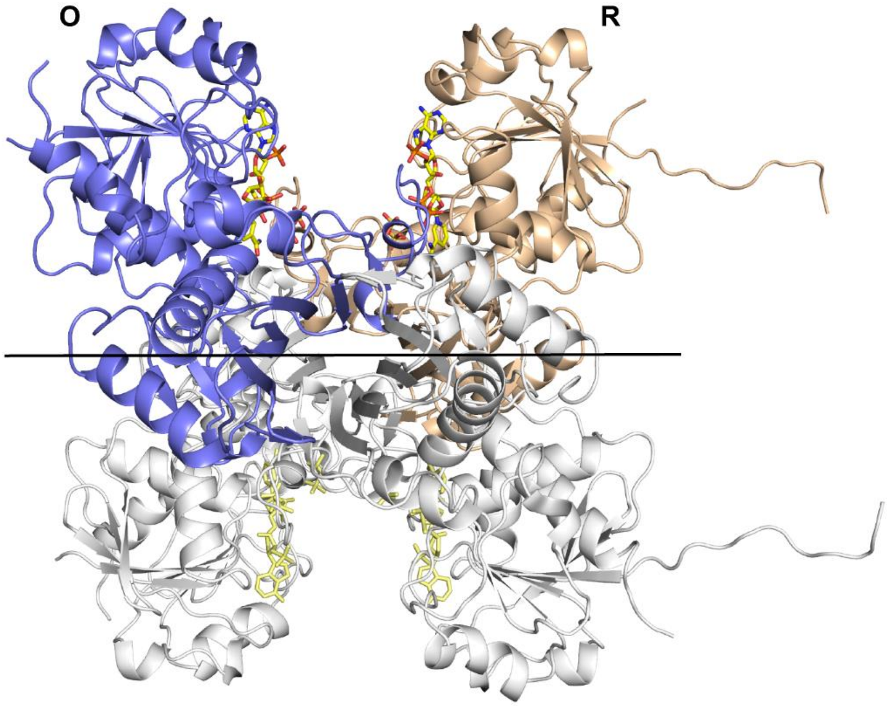


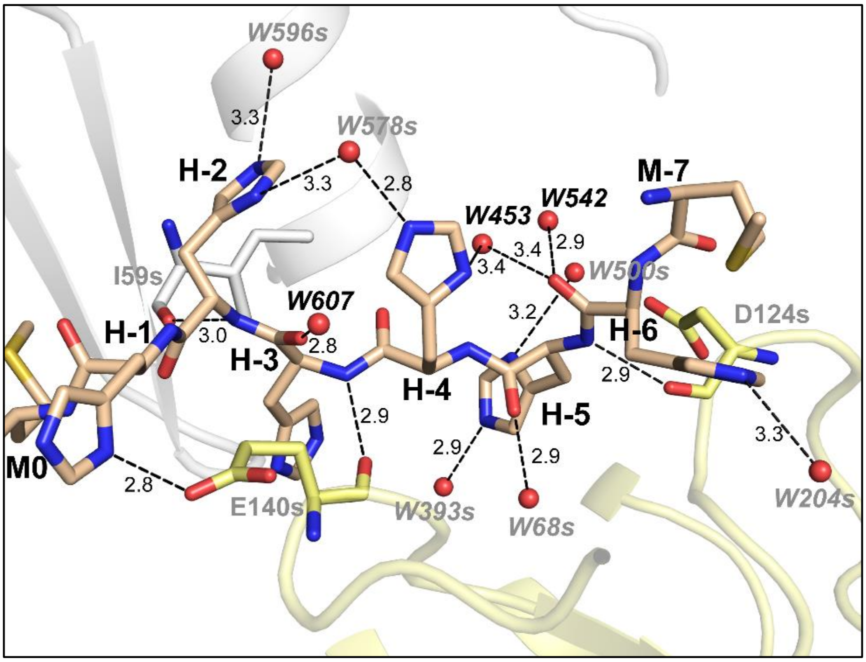


**Figure S6**. **Interactions of His-tag within NADP$^+$-CrGAPA crystal structure**. (a) Ribbon representation of the NADP$^+$-CrGAPA tetramer structure. The chain R shows at the N-terminal end the whole non-cleavable His-tag (MHHHHHHM). This portion disordered in chain O, breaks the 222 symmetry of the tetrameric structure which became a dimer of dimers (C2 symmetry). The symmetry molecular axis coincident with the 2-fold crystallographic axis is reported in black. The cofactor (NADP$^+$) bound to each monomer, is represented in stick. (b) Hydrogen bond interactions between the residues of His-tag (wheat ball and stick) and the residues or waters of symmetry related molecules (gray labels).

### Figure S6

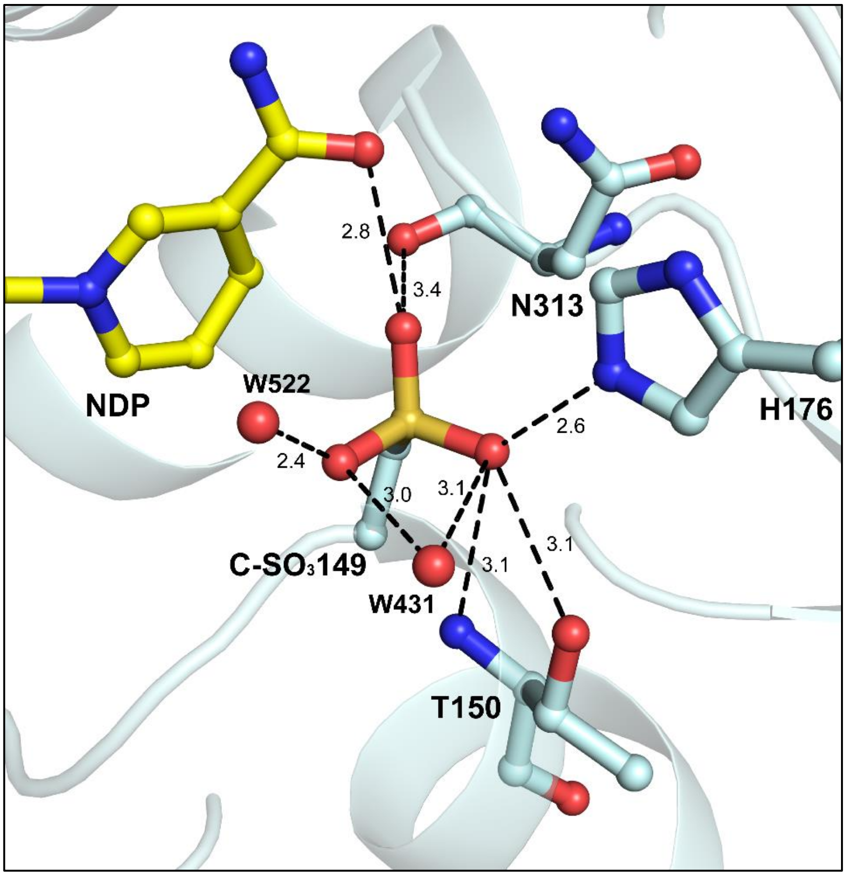


**Figure S7. Interactions of the catalytic sulfinyltated/sulfonylated Cys149 within oxidized NADP$^{+}$-CrGAPA crystal structure**. The sulphonate group (-$SO_3^-$) of Cys149 is stabilized by several interactions such as hydrogen bonds and electrostatic interactions (distance ≤ 3.5 Å) with residues, NADP$^{+}$, and water molecules.

### Figure S7

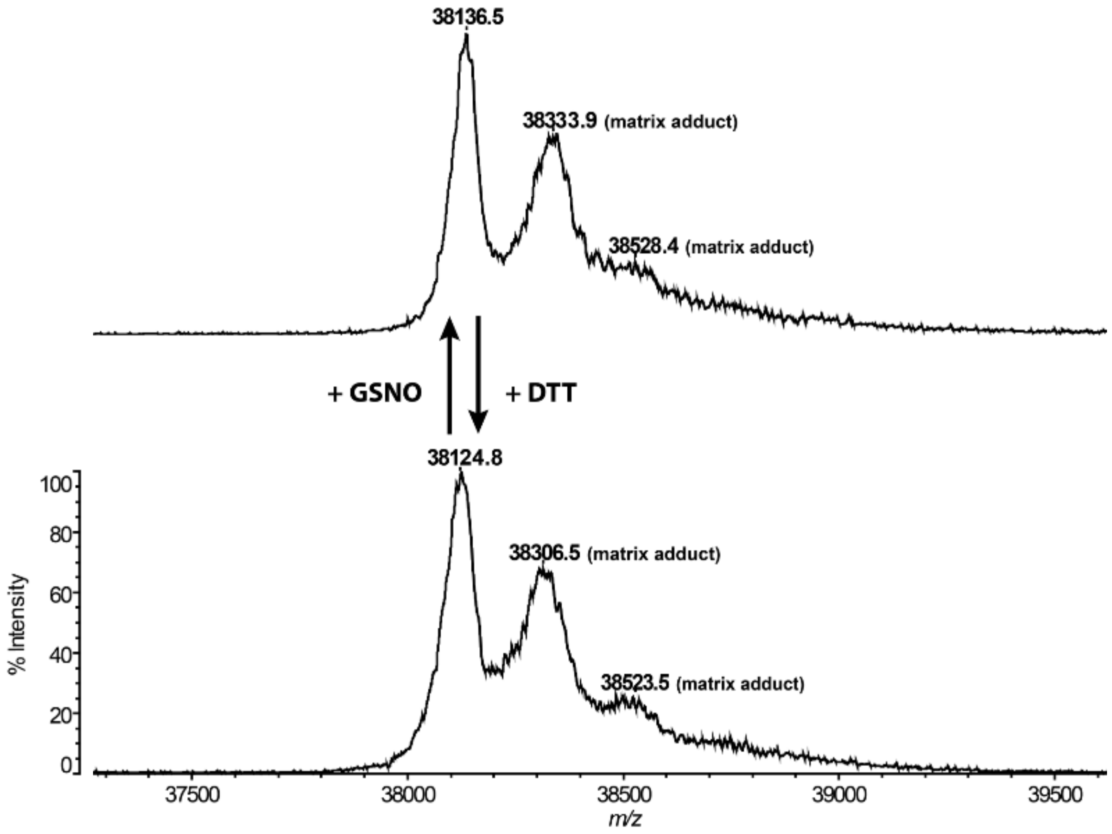


**Figure S7. MALDI-TOF mass spectrometry analysis of CrGAPA.** Mass spectra of CrGAPA treated with 2 mM GSNO for 30 min at room temperature were performed before (upper panel) and after (lower panel) a treatment in the presence of 20 mM DTT (30 min). The differences between mass peaks of GSNO-treated CrGAPA before (38,136.5 Da) and after DTT treatment (38,124.8 Da) are within the experimental error (0.1%) of the instrument. The peaks corresponding to CrGAPA with a sinapinic acid adduct are indicated.

### Figure S8

**a**

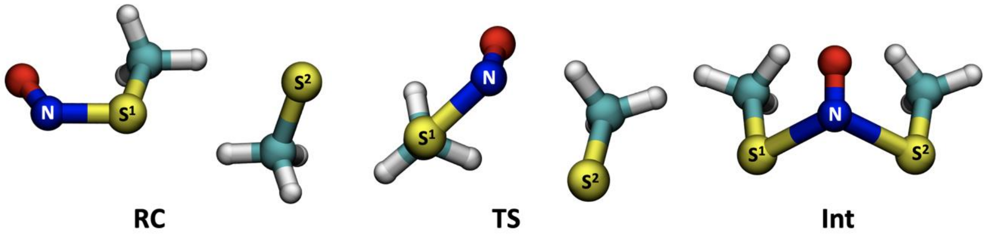


| | RC | TS | Int |
|---|---|---|---|
| d(N-S1) | 1.74 | 1.82 | 2.21 |
| d(N-S2) | 5.14 | 3.34 | 2.21 |

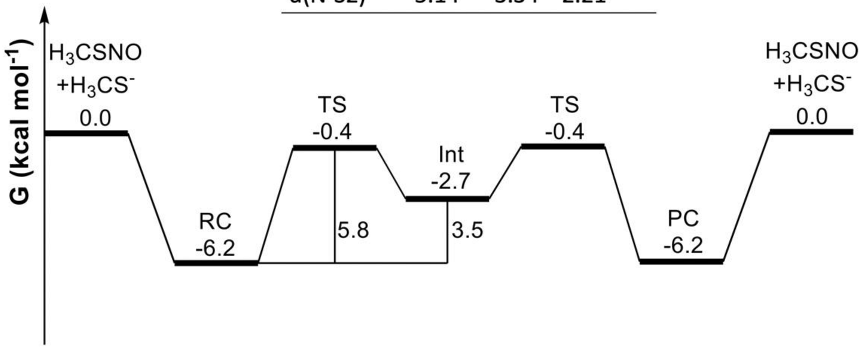


**b**

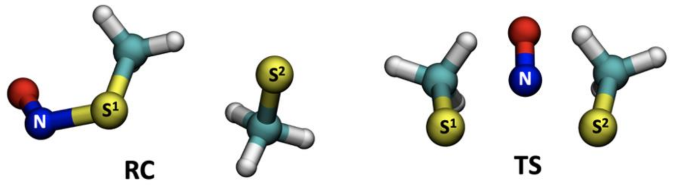


| | RC | TS |
|---|---|---|
| d(N-S1) | 1.74 | 2.14 |
| d(N-S2) | 5.37 | 2.34 |

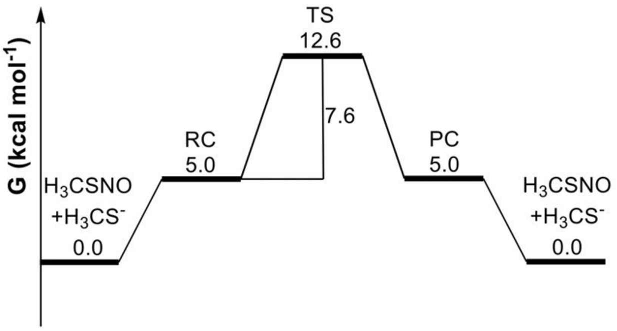


**Figure S8. Free energy surface (FES) for the trans-nitrosylation process in gas phase (a) and in water** (**b**). Activation barrier and reaction energies are expressed in kcal mol$^{-1}$. In the top of both panels, a 3D representation of the critical points (RC, reactive complex; TS, transition state; PC, product complex).

#### Table S1. Data collection and refinement statistics of CrGapA structures.

| | $NADP^+$ | $NAD^+$ | Oxidized |
|---|---|---|---|
| **Data collection** | | | |
| **Unit cell (Å)** | 141.98 148.51 74.06 90.00 90.00 90.00 | 143.34 147.10 75.16 90.00 90.00 90.00 | 142.40 147.33 74.76 90.00 90.00 90.00 |
| **Space group** | C222 | C222 | C222 |
| **Resolution range* (Å)** | 46.74 – 1.50 (1.53 – 1.50) | 60.64 – 2.20 (2.32 – 2.20) | 46.43 – 1.70 (1.73 – 1.70) |
| **Unique reflections*** | 124828 (6138) | 40233 (5844) | 84958 (4390) |
| **Completeness* (%)** | 100.0 (99.9) | 99.0 (99.4) | 98.4 (96.8) |
| **$R_{merge}$*** | 0.147 (0.794) | 0.203 (0.501) | 0.080 (1.164) |
| **$CC_{1/2}$*** | 0.995 (0.917) | 0.959 (0.846) | 0.999 (0.897) |
| **I/s(I) *** | 10.6 (3.0) | 3.6 (1.7) | 17.5 (1.8) |
| **Multiplicity*** | 13.1 (13.3) | 4.0 (4.2) | 13.0 (13.8) |
| **Refinement** | | | |
| **PDB ID** | 7ZQ3 | 7ZQK | 7ZQ4 |
| **Resolution range* (Å)** | 46.74 – 1.50 (1.52 – 1.50) | 42.42 – 2.20 (2.26 – 2.20) | 45.18 – 1.70 (1.72 -1.70) |
| **Reflection used** | 124793 (8049) | 38233 (2799) | 84144 (5344) |
| **$R/R_{free}$** | 0.187/0.205 | 0.249/0.267 | 0.2169/0.2289 |
| **rmsd from ideality (Å, °)** | 0.012, 1.550 | 0.008, 1.519 | 0.005, 0.945 |
| **N° atoms** | | | |
| **Non-hydrogen atoms** | 6151 | 5435 | 5973 |
| **Protein atoms** | 5307 | 5155 | 5260 |
| **Solvent molecules** | 713 | 162 | 572 |
| **Cofactor atoms** | 96 | 88 | 96 |
| **Hetero atoms** | 35 | 30 | 45 |
| **B value ($Å^2$)** | | | |
| **Mean** | 20.7 | 38.6 | 31.6 |
| **Wilson** | 13.8 | 31.5 | 24.9 |
| **Protein atoms** | 19.1 | 41.6 | 30.7 |
| **Cofactor atoms** | 15.4 | 46.7 | 35.4 |
| **Solvent molecules** | 32.8 | 36.6 | 38.4 |
| **Hetero atoms** | 39.1 | 61.9 | 43.8 |
| **Ramachandran plot (%)§** | | | |
| **Most favoured** | 95.6 | 95.7 | 96.1 |
| **Allowed** | 4.0 | 4.0 | 3.6 |
| **Disallowed** | 0.4 | 0.3 | 0.3 |

* Values in parentheses refer to the last resolution shell

§ As defined by MolProbity (42)
